# Antisense oligonucleotides targeting transcription termination windows disrupt mRNA 3’ end processing and decrease gene expression

**DOI:** 10.64898/2026.02.04.703595

**Authors:** Kinga Winczura, Lars Joenson, Erich Koller, Pawel Grzechnik, Łukasz Jan Kiełpiński

## Abstract

Antisense oligonucleotides (ASOs) are short, synthetic nucleic acids that bind to complementary RNA sequences and alter gene expression, making them versatile therapeutic agents. Here, we identify transcription termination windows of protein-coding genes as previously unrecognized targets for ASOs. We show that ASOs can act on nascent RNA synthesised downstream of the annotated genes, leading to a pronounced decrease in the corresponding mRNA levels. These downstream-of-gene ASOs (DG-ASOs) induce RNase H1-dependent cleavage, impairing mRNA 3’ end processing, directing the unprocessed mRNAs for exosome-dependent degradation and creating early entry points in the nascent RNA for the XRN2 exonuclease termination factor. Altogether, we show that termination windows are genuine ASO targets which can be exploited to suppress gene expression. Importantly, we also reveal an underappreciated source of off-target effects which may arise from ASOs binding downstream of genes. Our findings indicate the necessity to expand ASO design and off-target assessment guidelines to include termination window sequences, thereby improving therapeutic efficacy and safety.

## INTRODUCTION

Antisense oligonucleotides (ASOs) are short, single-stranded, modified nucleic acids that bind with high specificity to target cellular RNA via Watson-Crick-Franklin base pairing. ASO-RNA interaction may alter gene expression via multiple mechanisms, such as splicing modulation, miRNA inhibition or transcript degradation^1–3^. This versatility in regulating distinct mechanisms, coupled with favourable pharmacological properties^4^, makes ASOs a rapidly emerging class of therapeutic agents, with a number of approved drugs available on the market^2^.

One of the mechanisms used to reduce the expression level of a gene of interest harnesses the endogenous endonuclease RNase H1^5,6^. Upon ASO hybridization to the targeted RNA, the enzyme recognizes the DNA:RNA heteroduplex and hydrolyzes the RNA component^7^. The cleaved RNA is further recognised by the cellular surveillance machinery and degraded by exonucleases, leading to a decrease in steady-state mRNA and protein levels^8^. In this approach, a typical ASO features 8-12 deoxyribonucleotides flanked with three to five modified nucleotides on either side to protect the DNA gap from nucleolytic degradation and improve binding affinity^9,10^. For these so-called gapmer ASOs, the most common modifications in the flanking regions include locked nucleic acid (LNA), 2-O-methoxyethyl (2’-MOE) or constrained ethyl (cEt)^11–13^. Moreover, the DNA gap nucleotides can also be modified, for example, by the introduction of a single 2’-O-methyl (2’-OMe) modification, which has been described to significantly improve ASO properties and reduce cytotoxicity^14^.

After delivery into cells, ASOs become enriched in both nucleus and cytoplasm and may target nascent and steady-state RNA, respectively^15,16^. Nascent pre-mRNA is still attached to the transcribing RNA Polymerase II (Pol II) and contains introns which are co-transcriptionally removed by splicing^17–19^. Gapmer ASO-mediated RNase H1 cleavage in the introns may initiate premature transcription termination, contributing to silencing of the gene^15,16^. At the 3’ end, transcription continues beyond the annotated gene sequence into the so-called termination window^20^, potentially creating another region suitable for ASO binding. While ASOs targeting pre-mRNA intronic sequences are widely used^15,16^, it has not been explored whether they can act on nascent RNA transcribed downstream of genes within the transcription window and how this impacts gene expression.

Transcription termination coupled with mRNA 3’ end processing are essential processes in mRNA maturation, control of gene expression and establishing gene boundaries^21^. Pol II transcribing the 3’ end of the gene recruits the cleavage and polyadenylation factor (CPF), a large multiprotein complex orchestrating 3′ end processing coupled with transcription termination^21^. When the polyadenylation signal (PAS) appears in the nascent RNA, it is recognised by CPF and cleaved by the CPF endonuclease CPSF3^22,23^. The 3’ end of the pre-mRNA is polyadenylated by poly(A) polymerase to form a nuclear export-competent RNP to translocate to the cytoplasm^24^. If the maturation of the mRNA 3’ end fails, the transcript is recognised as faulty and directed for exonucleolytic degradation^25,26^. After reaching the PAS, Pol II stays bound to DNA and continues transcription within the termination window. This region in human cells has a median length of 3.3 kb but can extend to over 10 kb from the annotated end of the gene^20^. The Pol II complex transcribing the region downstream of PAS undergoes allosteric changes including dephosphorylation of elongation factors, leading to slowed transcription^21,27,28^. At the same time, the free RNA 5’ phosphate generated by CPSF3 cleavage over the PAS is recognized by the 5’-3’ exonuclease XRN2 which co-transcriptionally degrades nascent RNA, chasing the decelerating Pol II along the DNA template. The kinetic outpacing of transcription by XRN2-dependent degradation eventually results in the removal of Pol II from the DNA template ^27,28^.

In this study, we describe ASOs that act downstream of genes (DG-ASOs) by targeting nascent RNA synthesised within the termination windows where Pol II slowly transcribes prior to release from the DNA template. Treatment with DG-ASOs results in a reduction of mRNA levels and consequently, gene expression. DG-ASOs recruit RNase H1 which leads to co-transcriptional cleavage of the nascent RNA. This outcompetes cleavage performed by the CPF complex and affects mRNA 3’ end formation. The faulty mRNAs lacking correctly processed 3’ ends are degraded by the exosome, which results in a decrease of steady-state mRNAs and reduced protein levels. Additionally, DG-ASOs offer early entry points for XRN2, which accelerates degradation of nascent RNA within the termination windows. Finally, we demonstrate that a potential imperfect ASO off-target site located within a termination window can exhibit activity similar to on-target gene knockdown. Thus, our research expands the understanding of ASO mechanisms and highlights important aspects to consider when designing therapeutic ASOs.

## RESULTS

### DG-ASOs decrease steady-state mRNAs

Given that ASOs efficiently bind to and act on pre-mRNA^15,16,29^, we investigated whether they can target nascent RNA transcribed from regions located downstream of the PAS, within the transcription termination windows, and how this affects gene expression. In our study, we employed 16-nucleotide gapmer ASOs where the 10-nucleotide DNA gap was flanked with three LNA nucleotides on each side. We used them to target two unrelated genes, *CHMP1A* and *KPNB1,* that are characterised by different PAS usage, which allowed us to test ASOs in different genetic arrangements. *CHMP1A* uses a single PAS producing homogenous 3’ UTR transcripts, while *KPNB1* employs four different PAS motifs located within a 3 kb-long 3’ UTR, resulting in heterogeneous 3’ ends^30–32^. We designed two gapmer ASOs for each gene; one complementary to the annotated gene body (**g**ene **b**ody ASO; **GB-ASO**) and the other to a region located downstream of the last PAS (**d**ownstream of **g**ene ASO; **DG-ASO**) (Supplementary Table 1). For *CHMP1A*, the GB-ASO and DG-ASO targeted the first intron and the region located 132 nt downstream of the PAS, respectively (Fig. 1A). For *KPNB1*, the GB-ASO and DG-ASO targeted the last intron and the region located 120 nt downstream of the last PAS, respectively (Fig. 1B). Next, we transfected HEK293 cells with the ASOs targeting *CHMP1A* and *KPNB1* and a non-targeting negative control for 24 hours and tested the transcriptome by Bulk RNA Barcoding and sequencing (BRB-seq), an approach which produces 3’ cDNA libraries from polyadenylated RNA^33^. Analysis of the control samples confirmed that the sequences targeted by the DG-ASOs were positioned downstream of the last used PAS for both genes based on the most 3’ end peak on the BRB-seq tracks (Fig. 1A and B). To confirm that DG-ASOs targeted nascent RNA synthesised within the termination windows, we assessed active transcription downstream of *CHMP1A* and *KPNB1* genes using chromatin-associated RNA sequencing (chrRNA-seq). In this approach, nascent RNA transcripts are isolated from the chromatin fraction reflecting active transcription^34^. ChrRNA-seq disclosed the presence of nascent RNA that spanned at least ∼3 kb downstream of the *CHMP1A* PAS (Fig. 1A), as well as ∼4 kb downstream of the *KPNB1* last PAS (Fig. 1B). In both cases, the chrRNA reads overlapped with the DG-ASO target sequences, confirming the presence of nascent RNA available for ASO binding.

**Figure 1.**
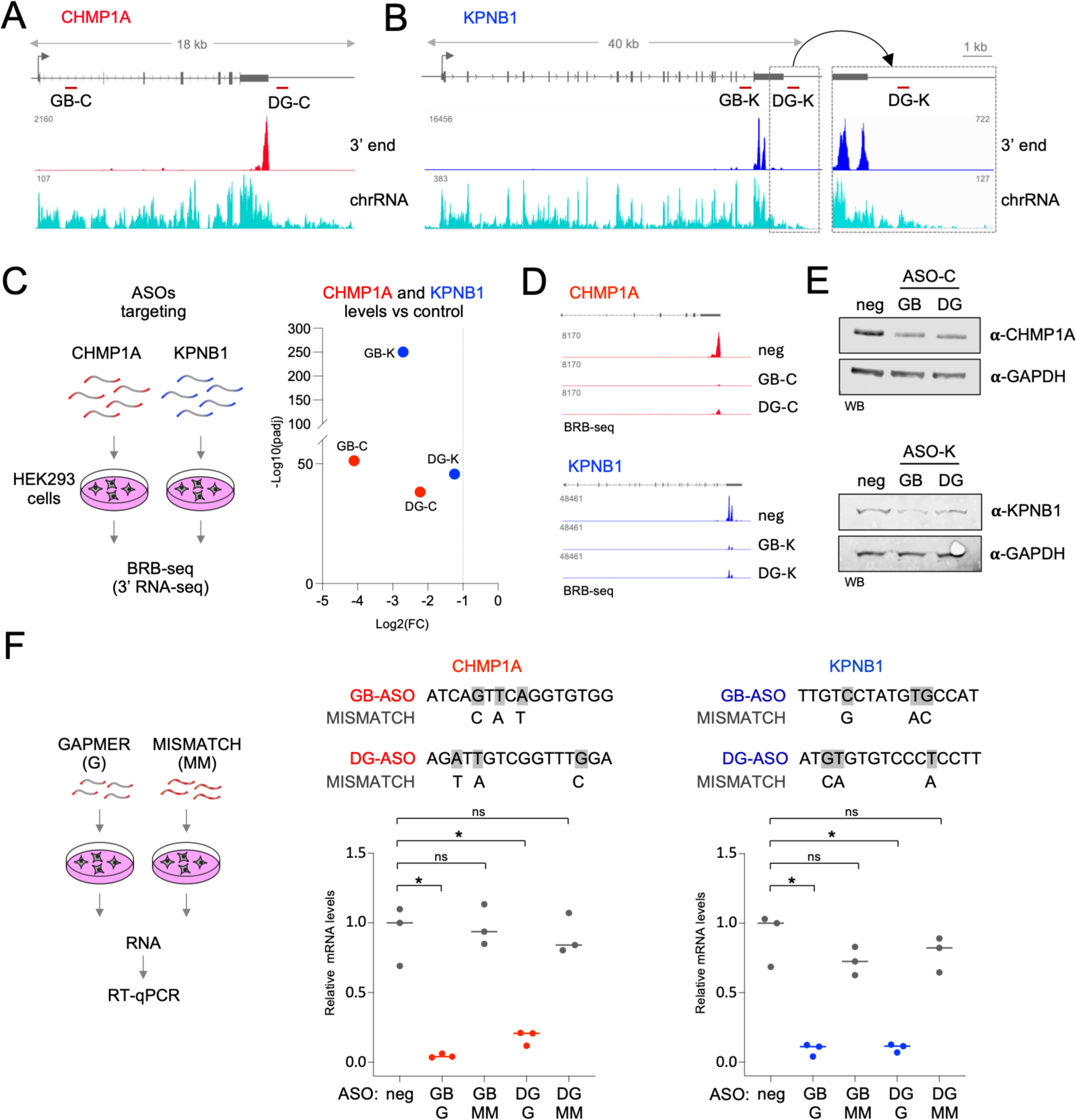
ASOs acting downstream of gene (DG-ASOs) affect mRNAs accumulation. **A-B.** The positions of ASOs targeting *CHMP1A* and *KPNB1* used in the study. *Top*, Genomic loci and the positions of ASOs relative to the annotated mRNA 3’ ends, with a zoom-in view for the 3’ end of the KPNB1 gene. *Middle,* IGV tracks showing *CHMP1A* (-C) and *KPNB1* (-K) PAS usage mapped by BRB-seq. *Bottom,* IGV tracks showing transcription indicated by chrRNA-seq reads. (GB – gene body, DG – downstream of gene) **C.** The levels of *CHMP1A* and *KPNB1* mRNAs upon GB- and DG-ASOs treatments quantified by BRB-seq. *Left*, a diagram showing the experimental approach. *Right*, a plot showing *CHMP1A* (-C) and *KPNB1* (-K) mRNAs fold change relative to the control. Differential gene expression analysis of three biological repeats. **D.** IGV tracks visualizing the sums of three replicates of BRB-seq reads. The scale is adjusted to the control track. (neg, negative control; treatment with a non-targeting ASO) **E.** CHMP1A and KPNB1 protein levels upon GB- and DG-ASOs treatments. Western blot analysis. GAPDH levels are shown as a loading control. **F.** The effect of mismatching ASOs on *CHMP1A* and *KPNB1* mRNAs. *Left*, a diagram showing the experimental approach. *Centre and right*, plots showing the levels of *CHMP1A* and *KPNB1* mRNAs after treatment with respective ASOs quantified by RT-qPCR. (GB – gene body; DG – downstream of gene; G – gapmer; MM – mismatch). Dots represent values of individual replicates. Horizontal lines represent the average of three biological replicates normalized to *GAPDH* mRNA and the median of the negative control (* p-val < 0.05, ns: p-val > 0.2, Student’s t-test).

Having established the specific transcriptional context for our ASOs, we analysed their impact on *CHMP1A* and *KPNB1* mRNA. Differential gene expression analysis of BRB-seq revealed that *CHMP1A* mRNA decreased by 94% (log_2_ FC = -4.1) upon treatment with GB-ASO and by 79% (log_2_ FC = -2.2) in cells transfected with DG-ASO (Fig. 1C). A similar effect was observed for KPNB1; GB-ASO decreased mRNA levels by 85% (log_2_ FC = -2.7) while DG-ASO by 58% (log_2_ FC = -1.2) (Fig. 1C). This was also evident from the BRB-seq tracks visualised in the Integrative Genome Viewer^35^ (Fig. 1D). We also tested the ASO-mediated knockdown efficiency on protein levels. Western blot analysis revealed that CHMP1A and KPNB1 protein levels in cells treated with GB- and DG-ASOs decreased, consistent with the decrease in mRNA levels measured by BRB-seq (Fig. 1E). To confirm the hybridization-dependent activity of GB- and DG-ASOs, we re-designed them to include three non-complementary nucleotides (Fig. 1F) based on the ASO experimental guidelines^36^. The cells were transfected with perfectly matching or mismatching GB- and DG-ASOs for 24 hours and the isolated RNA was tested by RT-qPCR. The levels of mRNAs were normalised to *GAPDH* mRNA and compared to the negative control. The mismatching ASOs did not affect mRNA levels compared with the perfectly matching ASOs which decreased the levels of *CHMP1A* and *KPNB1* mRNAs (Fig. 1F). Thus, we concluded that DG-ASOs interacted with RNA sequences downstream of the *CHMP1A* or *KPNB1* polyadenylation sites and this reduced the levels of the mRNAs that had been transcribed from the upstream located genes.

To further validate our observations, we examined DG-ASOs for two additional genes, proto-oncogenes *MYC* and *NET1*^37,38^ (Supplementary Table 1). We used our BRB-seq analysis in HEK293 cells to identify the most distal PAS for these genes and designed DG-ASOs complementary to the termination windows, 160 and 77 nt from the mature 3’ ends of *MYC* and *NET1*, respectively (Fig. 2A). The cells were transfected with DG-ASOs for 24 hours and the isolated RNA was assessed by RT-qPCR. *MYC* and *NET1* mRNA levels decreased after the treatment with their respective DG-ASOs, indicating that the ASO-dependent targeting of RNA within their termination windows affected the accumulation of steady-state transcripts (Fig. 2B). Finally, to confirm that the DG-ASOs downregulating effect is not cell line specific, we tested *CHMP1A, KPNB1, MYC* and *NET1* DG-ASOs in two additional cell types, neuroblastoma SH-SY5Y cells and HeLa cancer cells. These cells were transfected for 24 hours and the mRNA steady-state levels of *CHMP1A, KPNB1, MYC* and *NET1* were measured by RT-qPCR. Each DG-ASO reduced the level of corresponding mRNA in both cell lines (Fig. 2C). Note, that we used the same ASO delivery method that had been optimised for HEK293 cells, which may be less efficient in these cell types and potentially diminish ASO potency. Overall, our results indicate that the reduction in mRNA levels induced by DG-ASOs is a general phenomenon, and it depends on ASOs interacting with the RNA synthesised within the termination windows.

**Figure 2.**
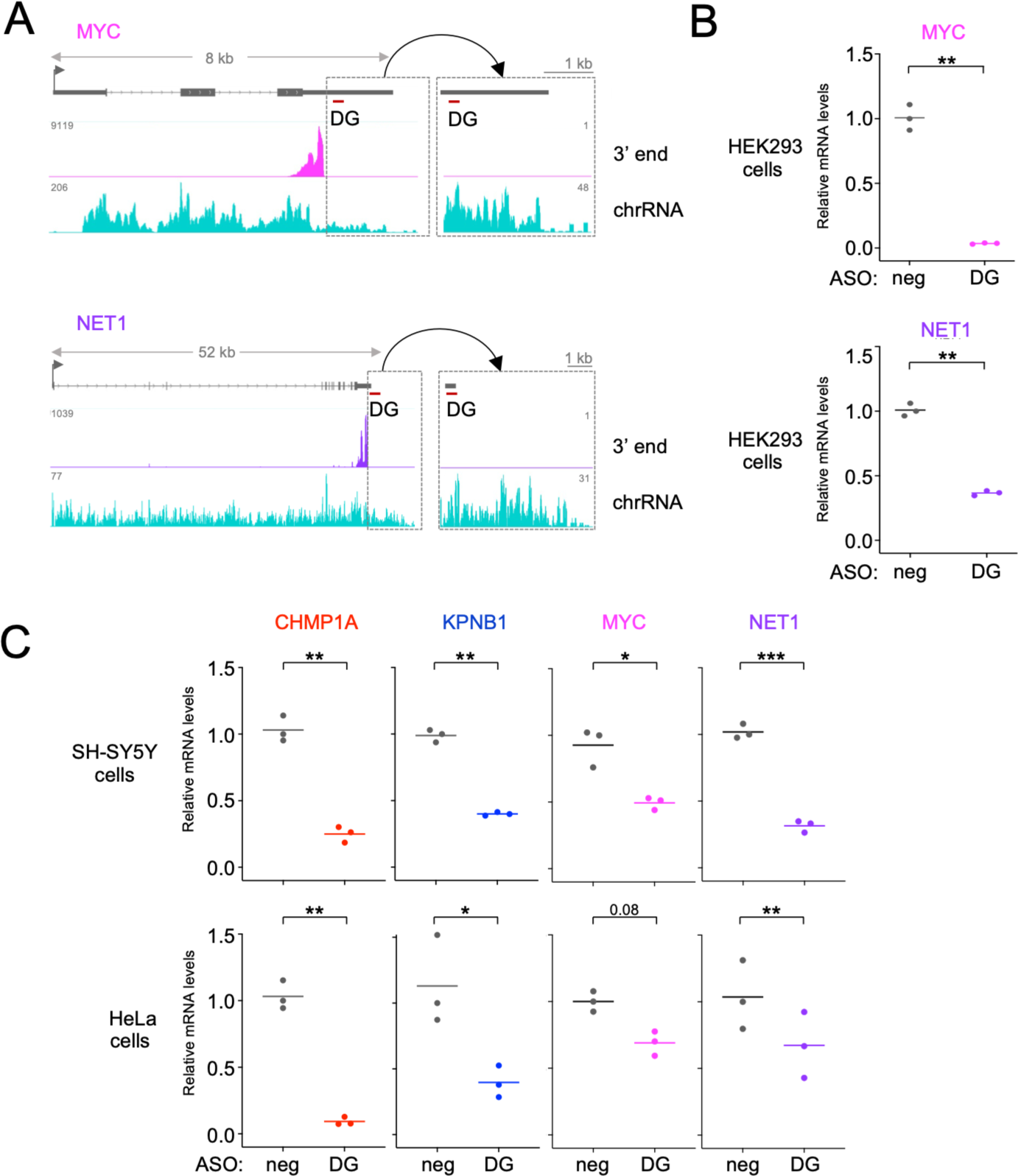
A universal effect of DG-ASOs on mRNA downregulation. **A**. The positions of ASOs targeting *MYC* and *NET1* mRNAs. Genomic loci and the positions of ASOs relative to the annotated mRNA 3’ ends are shown above the IGV tracks. The 3’ ends are mapped using BRB-seq (upper tracks). Active transcription is indicated by chrRNA-seq (lower tracks). 3’ ends are zoomed and magnified to confirm the lack of isoforms overlapping with DG-ASOs. (DG – downstream of gene) **B.** The levels of *MYC* and *NET1* mRNAs upon DG-ASOs treatments quantified by RT-qPCR. Dots represent values of individual replicates. Horizontal lines represent averages of three biological replicates normalized to *GAPDH* mRNA and median of negative control (** p-val < 0.01, Student’s t-test). (neg – negative control, DG – downstream of gene) **C.** The effect of DG-ASOs targeting *CHMP1A*, *KPNB1*, *MYC* and *NET1* in SH-SY5Y and HeLa cells measured by RT-qPCR. Dots represent values of individual replicates. Horizontal lines represent averages of three biological replicates normalized to *GAPDH* mRNA and median of negative control (*** p-val < 0.001; ** p-val < 0.01; * p-val < 0.05, Student’s t-test). (neg – negative control, DG – downstream of gene)

### RNase H is required for DG-ASO-dependent mRNA downregulation

The DG-ASOs used in our study were classical gapmers designed to form DNA:RNA hybrid substrates for RNase H cleavage, inducing RNA degradation. However, we could not exclude the possibility that the downregulation of mRNAs by DG-ASOs might be caused solely by their binding to the nascent RNA; therefore, we tested if this process is indeed dependent on RNase H.

First, we confirmed biochemical compatibility of DG-ASOs to induce RNase H cleavage *in vitro*. We designed T7 promoter-containing PCR primers flanking the regions complementary to either the *CHMP1A* or *KPNB1* DG-ASOs (approximately -100 and +100 bp from the binding sites) and used the resulting PCR products as templates for *in vitro* RNA synthesis. We hybridised the synthesised RNA fragments with their corresponding DG-ASOs and incubated with *E. coli* RNase H. As controls for each DG-ASO, we used a related ASO with mismatches, which has lowered binding affinity to the substrate, as well as a mixmer ASO, which contains locked nucleic acids instead of DNA nucleotides inserted throughout the central region (Fig. 3A and 3B). This modification prevents the mixmer from recruiting RNase H to the mixmer-RNA hybrid and consequently efficient RNA cleavage^7^. The products of the reactions were visualised on acrylamide gels. The most efficient cleavage activity was detected in samples incubated with gapmer DG-ASOs, indicating that the hybrid between the gapmer DG-ASO and RNA was effectively recognised by RNase H (Fig. 3B). Mixmer and mismatch ASOs caused only minor fragmentation of the RNA. Next, we employed *in cellulo* experimental approaches in HEK293 cells to demonstrate RNase H1 requirement for DG-ASO activity. We tested how gapmer ASOs impact *CHMP1A* and *KPNB1* mRNA levels upon RNase H1 knockdown. We depleted RNase H1 using siRNA for 24 hours, followed by transfection of *CHMP1A* and *KPNB1* GB- and DG-ASOs for another 24 hours. Western blot analysis confirmed the depletion of RNase H1 protein (Fig. 3C) and RT-qPCR analysis showed that the levels of *CHMP1A* and *KPNB1* mRNAs were rescued upon RNase H1 depletion (Fig. 3C). The effect for *KPNB1* was less pronounced likely reflecting poor co-transfection efficiency of the siRNA and both GB- and DG-ASO. Thus, we also tested the impact of ASO mixmers on *CHMP1A* and *KPNB1* mRNAs *in cellulo*. We transfected HEK293 cells with either gapmer or mixmer ASOs for 24 hours and assessed mRNA levels using RT-qPCR. None of the mixmer ASOs reduced *CHMP1A* or *KPNB1* mRNA levels (Fig. 3D), further confirming that RNase H1 cleavage over the ASO:RNA hybrid was required for DG-ASO-dependent mRNA downregulation.

**Figure 3.**
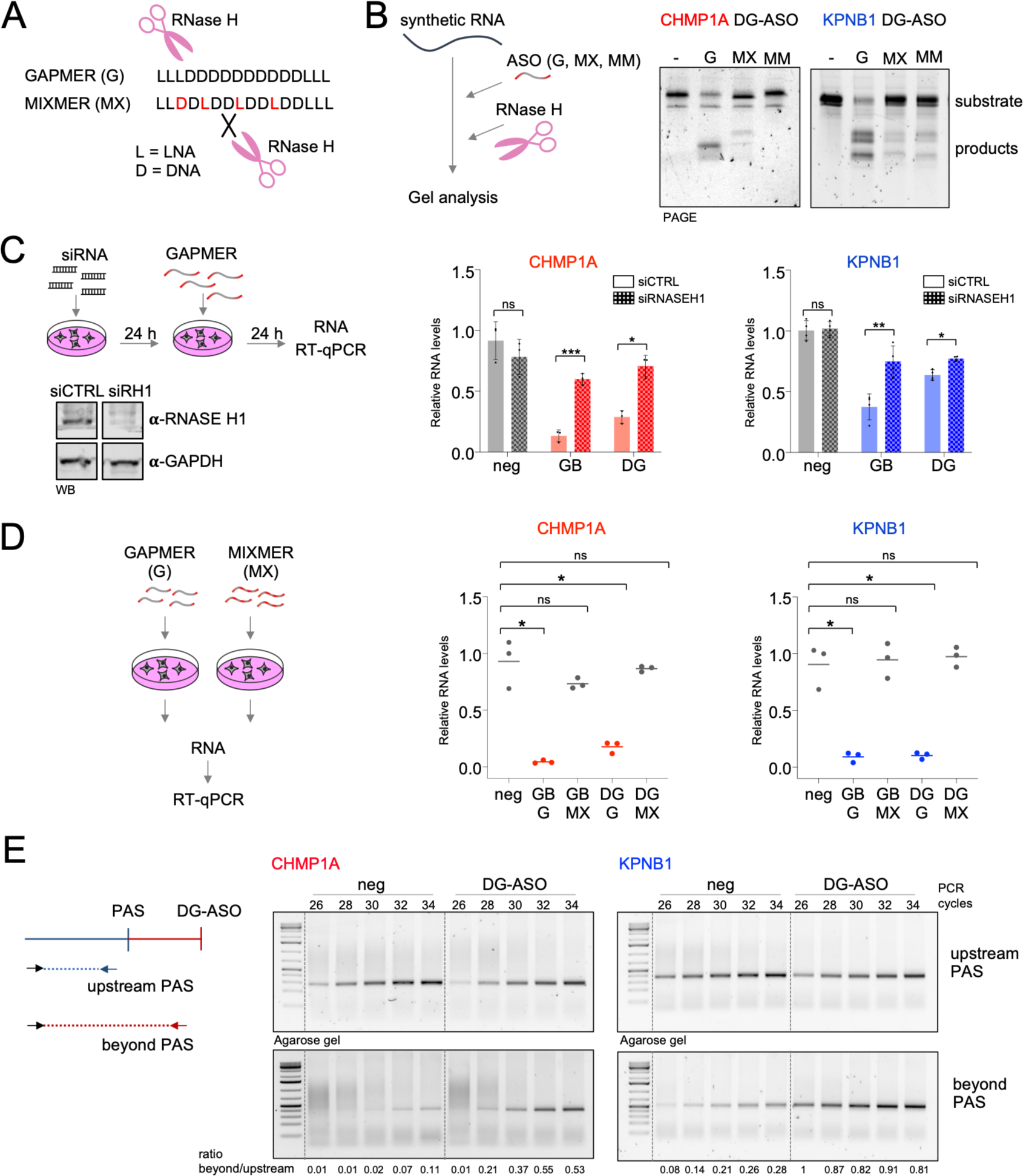
RNase H1 activity is necessary for the mRNA levels downregulation by DG-ASO. **A.** A diagram showing the composition of a mixmer ASO compared to a gapmer ASO. LNA or DNA nucleotides are labelled L or D, respectively. Red denotes change of the original nucleotide modification. **B.** *In vitro* processing of synthetic RNA targeted by DG-ASOs by recombinant RNase H. *Left*, a diagram presenting the experimental outline. Synthetic RNA containing the ASO binding site is incubated with different ASOs and RNase H. Reaction products are analyzed by PAGE. *Right*, denaturing PAGE analysis of cleavage products of *CHMP1A* or *KPNB1* synthetic RNA after incubation with respective gapmer (G), mixmer (MX) and mismatch (MM) ASOs. **C.** RNase H1 depletion impact on ASOs activity. *Left,* A diagram showing the experimental approach and, below, a western blot analysis of RNase H1 protein levels in control and RNase H1-depleted cells. GAPDH levels serve as a loading control. *Centre and right*, plots showing relative levels of *CHMP1A* and *KPNB1* mRNAs, measured by RT-qPCR, in cells treated with negative control (neg), gene body (GB) or downstream of gene (DG) ASOs following the control (siCTRL) or RNase H1 (siRNASEH1) protein depletion. Bars represent average RNA levels of three biological replicates normalized to *GAPDH* mRNA and the median of the negative control. Error bars represent standard deviation. Dots represent values of individual replicates (*** p-val < 0.001; ** p-val < 0.01; * p-val < 0.05; ns: p-val > 0.3, Student’s t-test). **D.** Mixmer ASO impact on *CHMP1A* and *KPNB1* mRNA levels. *Left,* a schematic of an experimental approach. *Right,* relative levels of *CHMP1A* and *KPNB1* mRNAs, measured by RT-qPCR, in cells treated with gapmer (G) or mixmer (MX) versions of gene body (GB) and downstream of gene (DG) ASOs. Dots represent values of individual replicates. Horizontal lines represent average RNA levels calculated from three biological replicates normalized to *GAPDH* mRNA and median of negative control (* p-val < 0.05, ns: p-val > 0.2, Student’s t-test). **E.** The accumulation of 3’ unprocessed *CHMP1A* and *KPNB1* mRNAs in cells treated with DG-ASOs measured by semi-qRT-PCR. *Left,* a schematic showing the localisation of PCR amplicons. *Right,* agarose gel analysis of PCR products. The number of PCR cycles for each reaction is shown above the gel image. The ratios of band intensities (beyond to upstream of PAS) were quantified by Fiji software and are shown below the gel image. (neg – negative control, DG – downstream of gene)

We hypothesised that the pre-mRNAs targeted by DG-ASO/RNase H1 may not yet have been cleaved and processed by CPF and so was continuous with the mRNA sequence. Thus, we asked how the DG-ASO-mediated cleavage affects CPF-dependent cleavage over the PAS. We designed RT-PCR primers to distinguish between the mature and the RNase H1-generated 3’ extended forms of *CHMP1A* and *KPNB1* mRNAs. A mutual forward primer was anchored in the 3’ UTR of each tested gene. Two reverse primers were designed; one upstream of the PAS to amplify both the mature CPF-cleaved and the extended forms and the second primer located beyond the PAS but upstream of the DG-ASO binding sites to amplify only the extended, RNase H1-cleaved forms (Fig. 3E). Since the fragments were unsuitable for qPCR analysis (due to their length ranging from 283 bp to 558 bp), we performed a semi-quantitative analysis by terminating the RT-PCR reaction every two cycles and analysing the products on agarose gels (Fig. 3E). For both genes, the levels of the amplicons located upstream of PAS were reduced (cycles 26 -30) in cells treated with DG-ASO, which is consistent with the observed DG-ASO-dependent *CHMP1A* and *KPNB1* mRNA downregulation. However, the products representing the uncleaved 3’ extended forms increased in the samples treated with DG-ASOs, indicating the elevated presence of unprocessed pre-mRNA compared to the control (the maximum difference in ratio beyond/upstream of PAS was 0.37 to 0.02 and 1 to 0.08 for *CHMP1A* and *KPNB1*, respectively). Taken together, DG-ASOs promoted RNase H1 cleavage of RNA downstream of PAS before CPF could cleaved the pre-mRNA over PAS. This resulted in the generation of 3’ extended faulty mRNAs and reduced the accumulation of steady-state mRNAs.

### DG-ASOs enhance transcription termination

Given that mRNA 3’ end processing occurs co-transcriptionally and is coupled with transcription termination, we decided to test how DG-ASOs impact these processes. We sequenced chromatin-associated RNA (chrRNA) obtained from cells treated with GB-and DG-ASOs. The quality and the nascence of the isolated RNA were assessed by measuring the ratio of chromatin-bound *NEAT1* lncRNA to predominantly cytoplasmic *GAPDH* mRNA. For all samples the ratio increased ∼20-40 times in the chromatin fraction which is indicative of robust nascent RNA enrichment^34^ (Fig. 4A). Differential gene expression analysis revealed that both GB- and DG-ASOs only moderately affected the levels of nascent RNA from *CHMP1A* or *KPNB1* loci (log2 FC *CHMP1A* GB-ASO = -1.5; log2 FC *CHMP1A* DG-ASO = -0.8; log2 FC *KPNB1* GB-ASO = -0.9; log2 FC *KPNB1* DG-ASO = -0.5) compared to the negative control ASO (Fig. 4B), indicating that the degradation happened mostly post-transcriptionally after RNA was released from the Pol II complex to the nucleoplasm. Only *CHMP1A* GB-ASO lowered nascent RNA levels below the threshold (log2 FC < -1), which can be explained by its co-transcriptional degradation by exonuclease XRN2^15,16^, since GB-ASO is located in the first intron, close to the 5’ end of *CHMP1A*. Note, that *KPNB1* GB-ASO was located towards the 3’ end of the gene and therefore XRN2-dependent degradation was not significant^15^.

**Figure 4.**
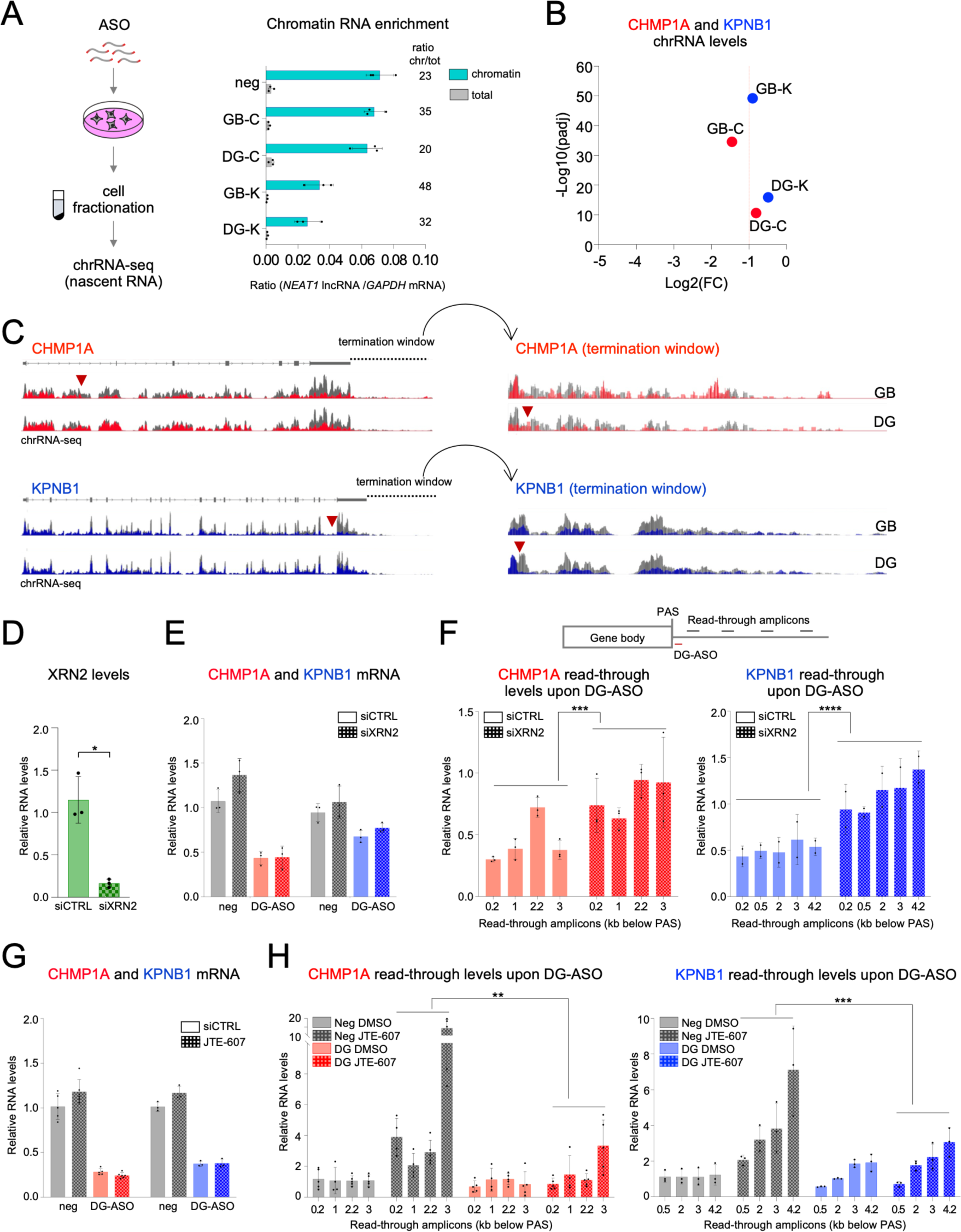
DG-ASOs impact on transcription termination. **A.** The experimental approach testing active transcription in the cells treated with ASOs. Plot on the right shows the ratio of *NEAT1* lncRNA to *GAPDH* mRNA in total and chromatin RNA fractions used as an indication of nascent RNA enrichment, RT-qPCR analysis. Bars represent the average of three biological replicates, error bars - standard deviation and dots represent values of individual replicates. (GB – gene body; DG – downstream of gene; C – *CHMP1A*; K – *KPNB1*; chrRNA-seq – chromatin-associated RNA-seq) **B.** The levels of *CHMP1A* and *KPNB1* nascent RNA upon GB- and DG-ASOs treatments quantified by chromatin-associated RNA-seq (chrRNA-seq). The plot shows *CHMP1A* and *KPNB1* nascent RNA foldchange relative to the control. Differential gene expression analysis of three biological repeats is shown. (GB – gene body; DG – downstream of gene; C – *CHMP1A*; K – *KPNB1*) **C.** chrRNA-seq reads visualised in Integrative Genome Viewer. Overlays of the average of three biological repeats for the negative control (grey) and ASO (red – *CHMP1A*, blue – *KPNB1*) treated cells are shown. The scale is adjusted to the control track. *Right,* zoom-in to the termination windows. Red arrowheads indicate ASO binding sites. (GB – gene body; DG – downstream of gene) **D.** RT-qPCR analysis showing the efficiency of *XRN2* knockdown. Bars represent average RNA levels of three biological replicates normalized to *GAPDH* mRNA and the median of the negative control. Error bars represent standard deviation. Dots represent values of individual replicates (* p-val < 0.05, Student’s t-test). **E.** The impact of *XRN2* depletion on DG-ASO mRNA downregulation activity. The plot shows relative levels of *CHMP1A* and *KPNB1* mRNAs in cells treated with negative control (neg) or downstream of gene (DG) ASOs following control (siCTRL) or XRN2 (siXRN2) depletion, measured by RT-qPCR. Bars represent the average RNA levels of three biological replicates normalized to *GAPDH* mRNA and the median of negative control. Error bars represent standard deviation. Dots represent values of individual replicates. **F.** Nascent RNA levels within the termination windows in cells treated with DG-ASOs and siXRN2. The amplicons used in the experimental approach are shown above the plot and the distance to PAS is indicated below each data point. The plot shows relative levels of nascent RNAs in cells treated with downstream of gene (DG) ASOs following control (siCTRL) or XRN2 (siXRN2) depletion, measured by RT-qPCR. Bars represent average RNA levels of three biological replicates normalized to *GAPDH* mRNA and the median of the negative control. Error bars represent standard deviation. Dots represent values of individual replicates (**** p-val < 0.0001; *** p-val < 0.001, Student’s t-test). **G.** The impact of DG-ASO-dependent mRNA downregulation on induced read-through transcription. The plot shows levels of *CHMP1A* and *KPNB1* mRNAs in cells treated with negative control (neg) or DG-ASOs followed by the inhibition of the CPF cleavage activity by JTE-607, measured by RT-qPCR. Bars represent average RNA levels of up to five biological replicates normalized to *GAPDH* mRNA and the median of the negative control. Error bars represent standard deviation. Dots represent values of individual replicates. **H.** The levels of induced read-through transcription upon treatment with DG-ASOs. The plots show the levels of *CHMP1A* and *KPNB1* nascent RNA within the transcription termination windows in cells treated with DG-ASOs followed by the inhibition of the CPF cleavage activity by JTE-607. The distance of each amplicon to the last PAS is indicated below the plots. The plots show relative levels of nascent RNAs compared to control. Bars represent average RNA levels of up to five biological replicates normalized to *GAPDH* mRNA and the median of the negative control. Error bars represent standard deviation. Dots represent values of individual replicates (*** p-val < 0.001; ** p-val < 0.01; Student’s t-test).

A detailed inspection of *CHMP1A* and *KPNB1* chrRNA-seq tracks (Fig. 4C) revealed two interesting observations. Firstly, sequencing read coverages in the 5’ regions of the genes were unaffected by any of the ASOs used. This indicates that DG-ASOs did not affect transcription initiation. Secondly, we observed reduced chrRNA-seq read coverage downstream of the ASO binding site, suggesting that co-transcriptional cleavage by RNase H may be followed by XRN2-mediated degradation as it has been reported previously for ASOs acting within gene bodies^15,16^. To confirm that DG-ASOs also generate entry points for XRN2, we depleted it using siRNA for 24 hours (Fig. 4D) followed by transfection of cells with DG-ASOs and RT-qPCR analyses. The XRN2 knockdown did not affect *CHMP1A* and *KPNB1* mRNA levels (Fig. 4E), indicating that this 5’-3’ exonuclease does not contribute to the degradation of the upstream RNase H cleavage products. Next, we designed amplicons spanning the transcription termination windows to measure the read-through nascent RNA as a proxy for Pol II transcription termination (Fig. 4F). We observed that the RNA levels downstream of DG-ASO binding sites were reduced by the treatment with DG-ASOs; however, this reduction was rescued to the control level upon XRN2 knockdown for both *CHMP1A* and *KPNB1* (Fig. 4F). This indicates that XRN2 was involved in the degradation of DG-ASOs 3’ cut-off products and that the XRN2-dependent degradation of nascent RNA within the *CHMP1A* and *KPNB1* termination windows was accelerated in cells treated with DG-ASOs.

Having previously demonstrated that DG-ASOs affected CPF cleavage over the PAS, we further investigated the interplay between the activity of the DG-ASOs and CPF-dependent transcription termination. For that, we uncoupled transcription termination from the CPF activity by using JTE-607, a prodrug^39^ for a small molecule inhibitor of the catalytic subunit of the CPF complex, CPSF3. JTE-607 directly binds to CPSF3, inhibits its endonucleolytic activity and blocks the release of newly synthesized pre-mRNAs^39–41^. First, we transfected cells with control and DG-ASOs for 24 hours to induce the cleavage of the nascent RNA by RNase H1 and then inhibited CPSF3 with the addition of JTE-607 for 2 hours. Incubation with JTE-607 did not change the levels of steady-state *CHMP1A* or *KPNB1* mRNAs (Fig. 4G). Next, we measured RNA downstream of PAS, within the termination windows. In control ASO samples, the amount of RNA from the termination regions of these genes highly increased after addition of JTE-607, indicative of read-through transcription consistent with the impairment of the cleavage over the PAS (Fig. 4H, grey checkered bars). However, when cells were transfected with DG-ASOs prior to incubation with JTE-607, the levels of the transcriptional read-through of *CHMP1A* and *KPNB1* did not increase (Fig. 4H, red and blue checkered bars). We concluded that DG-ASOs prevented the accumulation of read-through transcripts independently of the CPF activity. The inhibition of CPF cleavage led to a supply of the 3’ unprocessed substrates for DG-ASO/RNase H1, which in turn, created the entry for XRN2 exonuclease and consequently resulted in transcription termination.

### mRNAs targeted by DG-ASOs are degraded by surveillance mechanisms

Since DG-ASOs generate extended mRNAs with faulty 3’ ends, we next asked how these aberrant RNAs are degraded. Previous reports identified the RNA exosome complex as the degradation machinery responsible for the removal of upstream RNase H1 cleavage products^8^. To test if the exosome is involved in the degradation of 3’ extended mRNAs, we engineered a cell line with the catalytic subunit of the exosome, exoribonuclease DIS3^42^, tagged endogenously with FKBP12^F36V^ tag (degron-TAG, dTAG) using a CRISPR-Cas9 approach^43^. Induction of the proteasomal degradation of the tagged protein by the addition of the dTAG^V^1 molecule to the cell media led to virtually complete depletion of DIS3 within the first hour of treatment (Fig. 5A). The functional impairment of the complex was confirmed by the increased accumulation of non-coding RNAs called promoter-upstream transcripts (PROMPTs), known targets of the RNA exosome^44^ (Fig. 5A). To test whether DIS3 removes RNAs cleaved by ASO-dependent RNase H1 activity, we simultaneously treated cells with *CHMP1A* or *KPNB1* ASOs and dTAG^V^1 for 24 hours and measured levels of *CHMP1A* and *KPNB1* mRNAs by RT-qPCR. The DG-ASO-induced decrease in mRNA levels of both genes was rescued by DIS3 depletion, confirming the role for the exosome in the degradation of DG-ASO-generated aberrant mRNAs (Fig. 5B). Since *KPNB1* GB-ASO acts in the last intron, the reduced *KPNB1* mRNA level after treatment with this ASO was also rescued by the exosome inactivation, which is consistent with previous research reporting the role of the complex in the degradation of RNase H-cleaved mRNAs^8^. This was not observed however, in the case of *CHMP1A* GB-ASO, which targeted the 5’ end of the gene and would result in a short 5’ cut-off product (which we did not test for). We also examined the direct accumulation of the 3’ extended isoforms upon depletion of DIS3 by measuring the levels of RNA using amplicons located between the PAS and the DG-ASO targeting sites. We found 9.5- and 7.3-fold increases of the 3’ unprocessed *CHMP1A* and *KPNB1* mRNAs, respectively, in the cells treated with DG-ASOs and dTAG^V^1 (Fig. 5C), indicating that the exosome was involved in the degradation of these RNAs. Importantly, the accumulation of unprocessed RNAs was elevated by a synergistic effect of the double treatment with DG-ASOs and dTAG^V^1 (Fig. 5C). We also confirmed that the 3’ extended RNAs, which accumulated upon DIS3 depletion, were not cleaved over the PAS. We performed RT-PCR analysis using oligo (dT) primer for cDNA synthesis followed by PCR employing the forward primer, anchored in the 3’ UTR,and reverse primers, complementary to either a sequence upstream of the PAS or between the PAS and the DG-ASO binding sites. For both genes we observed the presence of the 3’ extended pre-mRNA forms after DG-ASOs treatment indicating that they had not been cleaved at the PAS. Their accumulation was further increased by the DIS3 depletion, confirming the role of the exosome in their degradation (Fig. 5D).

**Figure 5.**
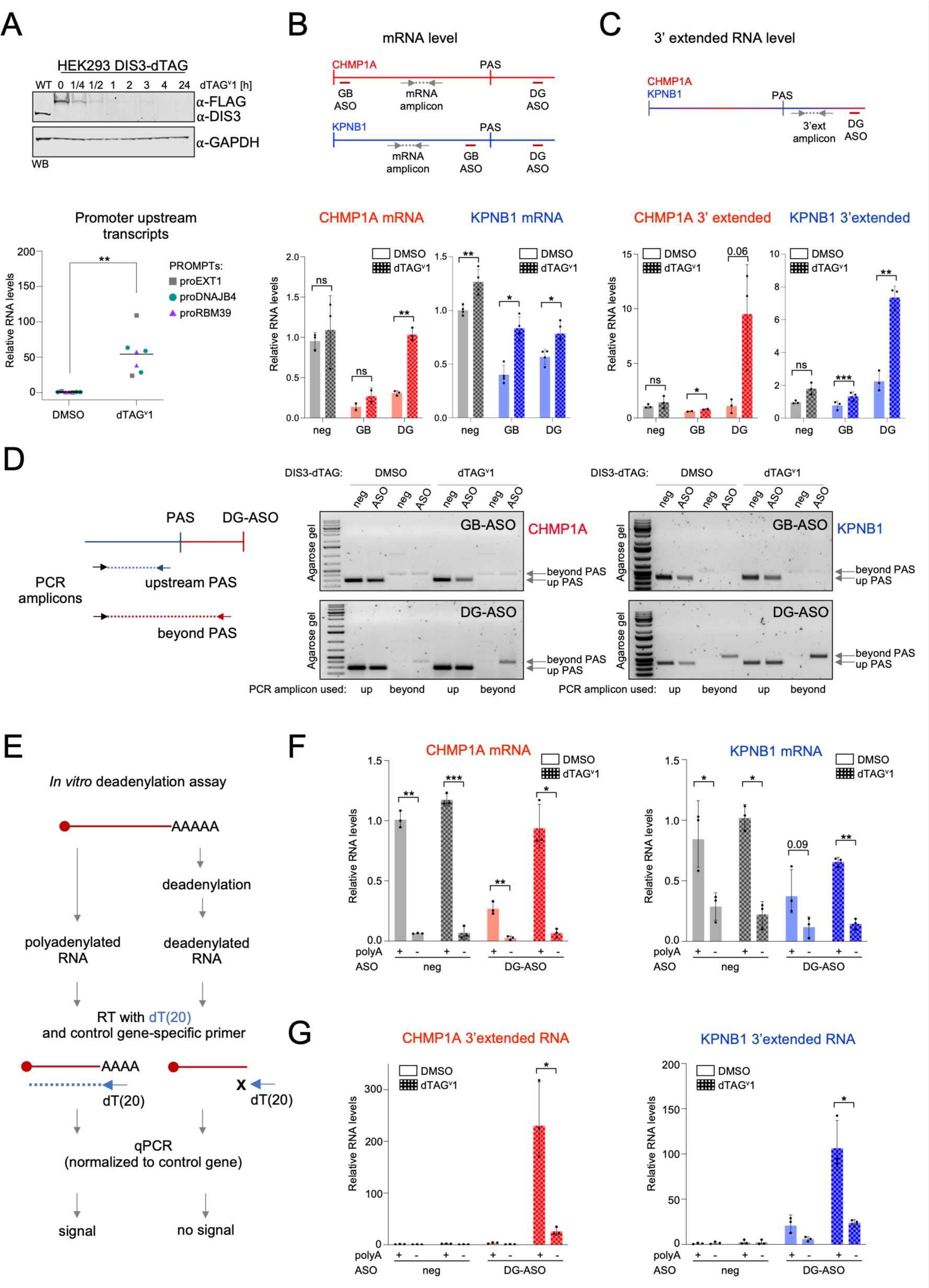
DG-ASO-generated aberrant mRNAs are polyadenylated and degraded by the exosome. **A.** Degron-tag (dTAG) induced depletion of DIS3. *Top*, western blot analysis showing the levels of DIS3-FKBP12^F36V^-3xFLAG in the cells incubated with dTAG^V^1 for the time indicated above the image. GAPDH is used as a loading control. *Bottom*, the accumulation of three different PROMPTs after 24 h incubation with DMSO or dTAG^V^1; RT-qPCR analysis normalized to *GAPDH* mRNA and median of the negative control. Horizontal lines represent averages of all values combined (** p-val < 0.01, Student’s t-test). **B - C.** Analysis of *CHMP1A* and *KPNB1* levels of mature (B) and 3’ extended (C) mRNAs in cells treated with DG-ASOs upon DIS3 depletion measured by RT-qPCR. The location of qPCR amplicons is shown above the plots. The plots show relative levels of mRNAs compared to the control. Bars represent average RNA levels of up to four biological replicates normalized to *GAPDH* mRNA and the median of the negative control. Error bars represent standard deviation. Dots represent values of individual replicates (*** p-val < 0.001; ** p-val < 0.01; * p-val < 0.05; ns: p-val > 0.05, Student’s t-test). **D.** The accumulation of 3’ unprocessed *CHMP1A* and *KPNB1* mRNAs in cells treated with DG-ASOs upon DIS3 depletion measured by semi-qRT-PCR. The location of PCR amplicons is shown on the left. The PCR reactions were terminated in late cycles (32-35), depending on the experimental set, to best visualise differences between the control and the treatment sample. The PCR products were analysed on agarose gels. (neg – negative control, GB – gene body, DG – downstream of gene) **E.** The experimental pipeline for the deadenylation assay. Oligo(dT) primer used for the removal of the poly(A) tail and subsequent reverse transcription is labelled dT(20). Deadenylated RNA levels are normalised to spike-in reverse transcription using control-gene (*NET1*) mRNA-specific primer. **F - G.** Analysis of the polyadenylation status of *CHMP1A* and *KPNB1* mature mRNAs (F) and DG-ASO-generated, 3’ extended forms (G) by RT-qPCR. The plots show relative levels of RNAs compared to the control. Bars represent average RNA levels of three biological replicates normalized to *NET1* mRNA and the median of the negative control. Error bars represent standard deviation. Dots represent values of individual replicates (*** p-val < 0.001; ** p-val < 0.01; * p-val < 0.05, Student’s t-test).

The DIS3-dTAG cell line allowed us to study the features of the 5’ RNA cleavage products by circumventing their immediate decay. Since the exosome can target polyadenylated and non-polyadenylated RNAs^45,46^, we aimed to investigate the polyadenylation status of the DG-ASO-generated, extended RNA forms. To that end, we isolated total RNA from cells treated with DG-ASOs in the background of DIS3 depletion and performed *in vitro* deadenylation. In this assay, we mixed oligo(dT) oligonucleotide with RNA and allowed for its hybridisation to mRNA poly(A) tails. The mix was then incubated with recombinant *E. coli* RNase H, which cleaved RNA within the DNA:RNA hybrid effectively removing the poly(A) tails. The polyadenylated (non-treated) and deadenylated RNAs were used as templates for reverse transcription with oligo(dT) primer. Since deadenylation disabled oligo(dT) priming, each reaction was spiked with a control mRNA-specific primer recognising the exonic sequence of *NET1* used for normalisation (Fig. 5E). As expected, the mature *CHMP1A* and *KPNB1* mRNAs were detected in polyadenylated fraction and their levels greatly decreased in deadenylated samples confirming the efficiency of deadenylation (Fig. 5F). Remarkably, the DG-ASO-generated 3’ extended *CHMP1A* and *KPNB1* mRNAs were also efficiently reverse-transcribed using oligo(dT) primer. They increased by more than 200-fold for *CHMP1A* and 100-fold for *KPNB1* upon DIS3 depletion compared to control. However, the deadenylation treatment reduced their signals by ∼9-fold and ∼4.5-fold, respectively (Fig. 5G). This strongly indicates that these 3’ extended transcripts resulting from DG-ASO/RNase H-dependent cleavage downstream of PAS were also polyadenylated.

### Termination windows represent a source of ASO off-target activity

We demonstrated that sequences downstream of genes, namely transcription termination windows, are viable ASO targets. Consequently, these sequences may contribute to off-target responses as any unwanted hybridisation in these regions could affect gene expression. Therefore, termination windows should be considered in the guidelines for ASO design. As a proof of principle, we investigated changes in mRNA levels in THP1 cells treated with a promiscuous ASO (pASO) (Supplementary Table 1), which was designed to target cancer-related hypoxia inducible factor 1 alpha *HIF1A* and used previously in multiple studies^47–51^. This ASO is also perfectly complementary to nine other mRNAs (Fig. 6A) and we found that it has an imperfect but favourable match (containing one terminal mismatching nucleotide^52^) to a sequence located outside of the annotated gene sequence of *CHMP1A*, 236 nucleotides downstream of the PAS (Fig. 6B).

**Figure 6.**
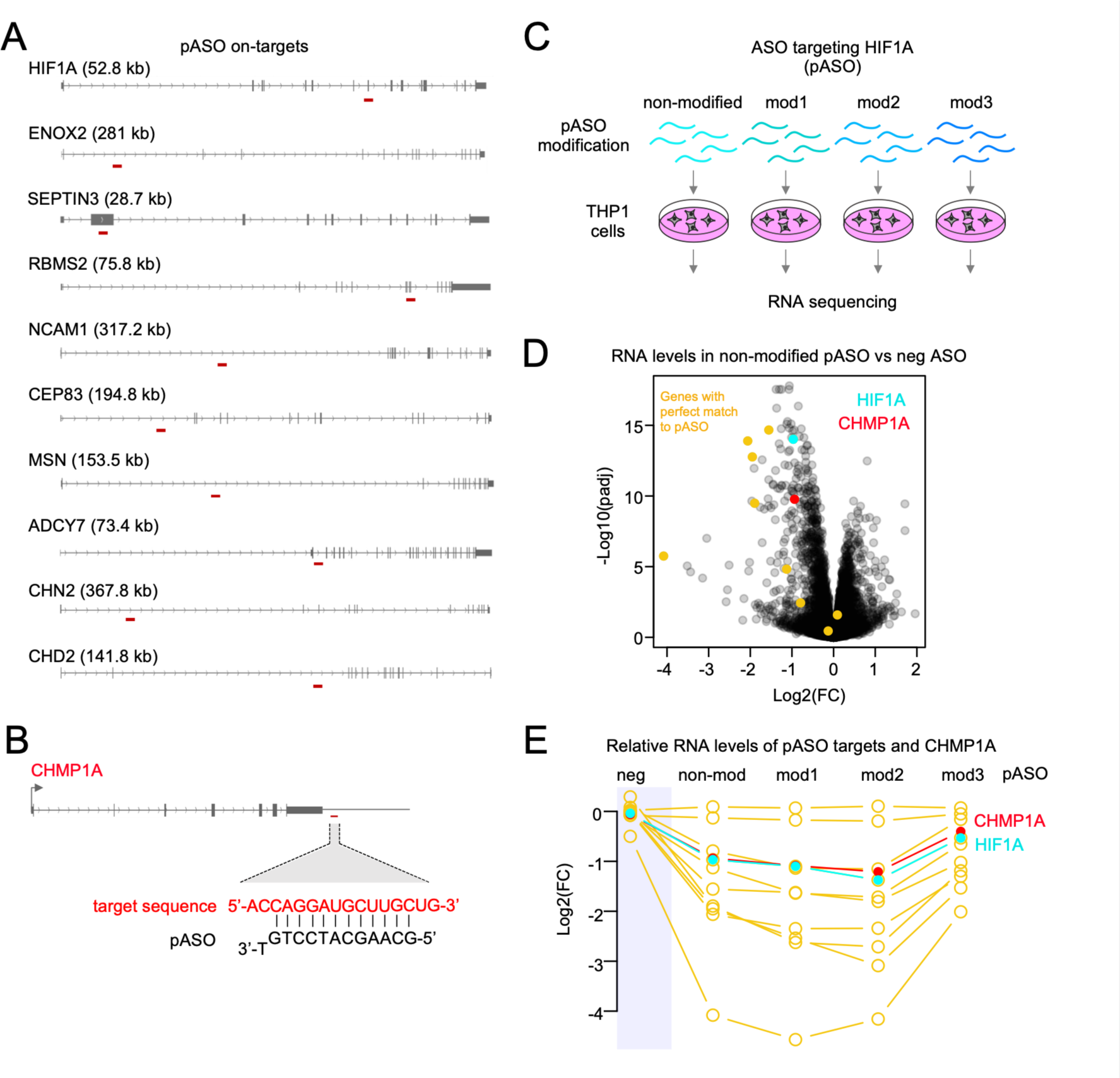
Off-target DG-ASO mimics the target ASO action. **A.** Genes targeted by pASO. The binding of pASO is shown below gene diagrams. **B.** The *CHMP1A* locus with a zoom into the pASO binding site in the region downstream of the gene. The RNA sequence complementary to pASO is shown in red. **C.** A diagram showing the experimental approach. Mod1-3 represent different 2’-O-methylated (2’-OMe) forms of pASO. **D.** Differentially expressed genes in THP1 cells treated with pASO, relative to control cells; RNA-seq analysis. mRNAs with complementary sequence perfectly matching to pASO are labelled in yellow, *CHMP1A* mRNA is labelled in red, *HIF1A* mRNA is labelled in cyan. **E.** The levels of pASO on-target mRNAs (yellow), *HIF1A* (cyan) and *CHMP1A* (red) mRNAs in cells treated with differentially 2’-OMe-modified pASOs. The negative control is on a blue background. (non-mod – non-modified pASO; mod1-mod3 – differentially modified pASO)

Leveraging the promiscuous character of pASO, we compared its impact on on-target mRNAs versus its off-target effect in the *CHMP1A* termination window. We tested if the addition of a 2’-O-methylation (2’-OMe) at various positions of the pASO central gap, a modification used for improving the therapeutic index of oligonucleotides^14^, could affect pASO off-target effect on *CHMP1A*. We treated THP1 cells with non-modified and three variants of pASO (Fig. 6C), each containing a 2’-OMe modification on a different nucleotide (Supplementary Table 1) and employed RNA-seq analysis to detect changes in mRNA levels compared to non-treated control cells. Differential gene expression analysis revealed that treatment with non-modified pASO resulted in decreased levels of seven target mRNAs (Fig. 6D). Strikingly, *CHMP1A* mRNA was downregulated similarly to other on-target genes, including *HIF1A* (log2 FC -0.92 vs -0.96, respectively), confirming that off-target interaction in the termination window meaningfully affected the accumulation of steady-state mRNA. The analysis of RNA from cells treated with modified pASOs revealed that the mRNA levels of *CHMP1A* fluctuated in response to pASO methylations in the same way as of the on-target genes (Fig. 6E). This indicates that the effect of ASO acting within the termination window did not differ from on-targeting activities in the gene bodies, confirming *CHMP1A* as a bona fide hybridization-based off-target of pASO. Overall, our results demonstrate the critical importance of considering termination windows when designing ASOs.

## DISCUSSION

ASOs are considered one of the cornerstones of personalized medicine and a number of them are already in therapeutic use or in clinical trials^2^. However, there are still significant gaps in our knowledge about their mechanism of action. Here, we report that ASOs can act via an RNase H1-dependent mechanism downstream of the annotated gene sequences, within the gene transcription termination regions. Such downstream of gene ASOs (DG-ASOs) trigger co-transcriptional cleavage of nascent RNA, affecting correct mRNA 3’ end processing and leading to the degradation of 3’ extended transcripts (Fig. 7).

**Figure 7.**
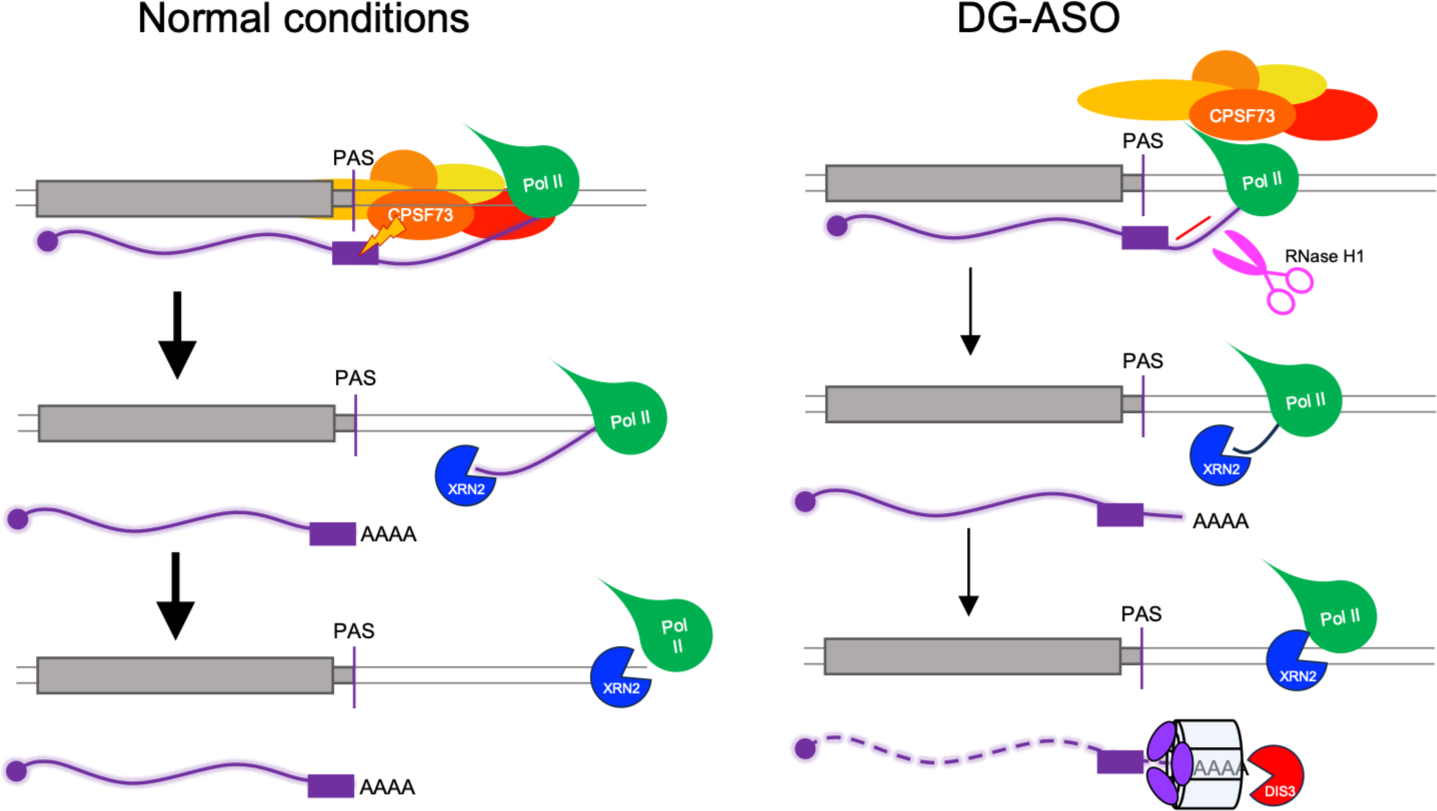
A model showing the mechanism of DG-ASO action. In normal conditions, nascent RNA is cleaved by CPF over the PAS sequence and the newly created 5’ end is attacked by exonuclease XRN2, which chases transcribing Pol II, effectively terminating transcription. In the cells treated with DG-ASO, RNase H1-dependent co-transcriptional cleavage outcompetes CPF inhibiting its ability to cleave at the PAS. As a result, the entry for XRN2 is created earlier, which may contribute to the faster transcription termination. The faulty, 3’ unprocessed pre-mRNA is polyadenylated and degraded by the exosome.

### Implications for the ASO design guidelines

The key inference of this study is the expansion of the sequence span targetable by RNase H1-recruiting ASOs to regions located downstream of mRNA polyadenylation sites. Our discovery should be considered as having both beneficial and detrimental consequences for ASO usage and as such, it implies the need to revise general guidelines for ASO drug design.

Firstly, our results indicate the necessity to update the off-target considerations to include RNA synthesized in the termination windows. Unintended hybridization events to the off-target RNAs are a critical safety concern for therapeutic ASOs, as they may introduce undesirable alterations in gene expression^52^. To avoid off-target effects, current guidelines recommend aligning ASO sequences with transcriptome databases containing sequences of mature mRNA, non-processed pre-mRNA and noncoding RNAs in order to select sequences exclusively matching the target gene^52^. Our results clearly indicate that sequences encompassing termination windows, which are not directly annotated in databases and so, not included in the current guidelines, should also be assessed for potential off-target binding of RNase H1 recruiting ASOs, as binding in those regions can trigger gene knockdown.

On the other hand, the addition of the termination windows to the pool of targetable sequences increases the opportunity to design efficient gene-specific ASOs. This can apply to virtually all Pol II transcription units but may be particularly useful for knocking down shorter genes if finding highly potent ASOs within gene bodies proves challenging. Using DG-ASOs also offers a unique opportunity to target intron-less genes when RNA downregulation early in biogenesis is preferred, e.g. to prevent the export of processed mRNA to the cytoplasm. Employing the termination windows may also increase the opportunity to find suitable targetable sequences containing single nucleotide variants (SNVs). The mutated mRNAs could be removed in an allele-specific manner, making the approach applicable to personalised medicine and the treatment of rare diseases^53–56^. Moreover, the use of DG-ASOs may be beneficial when targeting highly structured non-coding RNAs. Although successful ASO-dependent reductions for such ncRNAs including small nucleolar RNAs (snoRNAs), *MALAT1* and *NEAT1* have been reported^57–59^, targeting other structured transcripts may still pose a problem. Three-dimensional RNA motifs and structures can interfere with the ASO:RNA hybridization and contribute to ASOs inefficiency and/or cytotoxicity^60–62^. Thus, targeting the nascent RNA synthesised in the termination window may provide alternative options for such genes.

In our study, we used DG-ASOs located at most 236 nt downstream of the most distal PAS (off-target pASO, Fig. 6B). However, in the case of *KPNB1* gene, DG-ASO was positioned respectively 2 and 2.5 kb downstream of the main two proximal PASs and still exhibited satisfactory activity. Considering the wide variation in the length of termination windows^20^, we expect that precise boundaries and distances from the polyadenylation sites at which DG-ASOs remain effective will need to be individually established for specific genes, cell types, tissues and organisms most likely depending on the strength of the poly(A) signal^63^.

### Mechanistic insights into DG-ASO/RNase H1-dependent RNA degradation

RNase H1 is involved in essential cellular processes^64^. This conserved enzyme recognises DNA:RNA hybrids and cleaves their RNA component, and its activity is required in the removal of Okazaki fragments during replication and resolving of three strand structures called R-loops which form during transcription^64–66^. The RNase H-dependent cleavage removes the DNA:RNA hybrid, however our knowledge of how the RNA molecule is further processed is limited.

DG-ASO-dependent RNA cleavage misaligns the correct 3’ end, marking the mRNA as faulty, which consequently leads to its degradation and removal from the cell. The upstream product of the RNase H cleavage within the gene body is degraded by the exosome, the major 3’ to 5’ degradation machinery^8^. Our data shows that the depletion of DIS3, a catalytic subunit of the exosome, results in the accumulation of DG-ASO-generated 3’ extended and polyadenylated mRNA. This is consistent with the fact that exosome targets polyadenylated RNAs^45,46^; however, it is not clear if polyadenylation is required for the degradation of DG-ASO-generated mRNAs. Specific recruitment of the exosome to RNA is provided by two adaptor complexes; the nuclear exosome targeting (NEXT) complex targets non-polyadenylated shorter transcripts, whilst the poly(A) tail exosome targeting (PAXT) complex targets longer polyadenylated transcripts^67,68^. Despite this distinction, NEXT targets can be post-transcriptionally adenylated or uridylated and transferred to the PAXT-mediated decay pathway if NEXT or the exosome components are functionally deficient^46^. We show that the upstream products of DG-ASO-dependent RNase H1 cleavage are polyadenylated; however, it is also possible that they are polyadenylated upon DIS3 depletion due to the lack of degradation process. It remains to be determined what the length and composition of the tails are, and whether this polyadenylation is performed by the canonical CPF poly(A) polymerase due to the spatial proximity of the DG-ASO binding sites to the PAS, or by surveillance-associated polymerases such as TENT4A/B^69^. Overall, we show that DG-ASO-targeted, RNase H1-cleaved nascent RNAs are polyadenylated, which indicates the existence of the polyadenylation pathway for RNase H products.

We also show that downstream products of DG-ASO-mediated RNase H1 cleavage are removed by XRN2 exoribonuclease, as previously reported for ASOs targeting exonic and intronic gene sequences^15,16,28^. DG-ASO treatment results in the synthesis of 3’-extended RNAs, which are not cleaved over the PAS sequence. This indicates that RNase H1 can access and cleave nascent RNA earlier than CPF, although the DG-ASO binding and cleavage site is synthesized later than that of CPF, inferring competition between these two endoribonucleases. The binding of ASO to nascent RNA has been well documented and used to create an entry in RNA intronic sequences or to alter mRNA splicing^15,16,29,70^. Now, we show that the cleavage of the ASO:RNA hybrid by RNase H1 is rapid and suggest that it may kinetically outcompete physiological 3’ end formation. The fast action of RNase H1 may be associated with its function in removing R-loops. These structures are not only the source of DNA mutations but also block transcription and replication and, therefore, must be rapidly resolved^65^. The RNase H1/ASO-dependent co-transcriptional cleavage in the last exon upstream of PAS can also facilitate termination^28^. The 3’ end processing defect introduced by DG-ASOs indicates that CPF may not cleave pre-mRNA immediately after the PAS becomes available in the nascent RNA, especially as the regulatory elements may be located up to 100 nt downstream of the PAS^25,71,72^. Since the kinetics of the CPF-dependent cleavage is rapid once the complex is constituted and bound to the RNA^73–75^, the assembly of the complex on the chromatin or recognition of the PAS may be the limiting factors^63^, resulting in slower kinetics of the 3’ end formation than the recognition of the ASO:RNA by RNase H1. The presence of ASOs on the nascent RNA is not sufficient to affect the function of CPF, as the DG-ASO mixmers, which bind to target RNA but cannot recruit RNase H, did not affect mRNA levels. The rapid introduction of the cleavage by RNase H/DG-ASO within the termination window provides earlier access for the crucial termination factor, XRN2 exonuclease. Consistently, we show that DG-ASO treatment results in reduced nascent RNA within the termination window, indicating accelerated termination. We hypothesise that DG-ASO treatment does not affect the formation of the CPF complex which remains intact on the chromatin. By doing so, it enforces allosteric changes in the transcribing complex that affect the Pol II transcription rate to provide necessary conditions for XRN2 to act in transcription termination.

Overall, we describe a new ASO-mediated mechanism for gene expression reduction that expands the targeting opportunities and has important implications for considering off-target effects. DG-ASOs acting within the termination windows affect mRNA 3’ end processing and termination and, as a result, decrease mRNAs and, consequently, protein levels. Our results shed new light on RNase H1 activity *in vivo*, mRNA 3’ end processing and transcription termination, and we expect DG-ASOs to become not only expanded therapeutic candidates but also novel precision tools to dissect the kinetics of transcription termination.

## Acknowledgments

This work was funded by F. Hoffmann-La Roche Ltd, Switzerland. We thank Alan Whitmarsh, Cathy Tournier and Matthias Soller for critical reading of the manuscript and the UoM Genomics Technologies facility and Alithea for the support in RNA sequencing.

## Contribution

LJK conceived the study. PG, KW and LJK designed the experiments, analysed and interpreted the data. LJ carried out experiments in THP1 cells. KW and PG performed the rest of the experimental approaches (90% and 10%, contribution, respectively). EK contributed to data analysis and interpretation. PG and KW wrote the manuscript with LJK assistance.

## METHODS

### Cell culture media

HEK293, HeLa, SH-SY5Y and THP1 cells were maintained under standard culture conditions in a humidified incubator at 37 °C with 5% CO₂. HEK293 and HeLa cells were cultured in Dulbecco’s Modified Eagle Medium (DMEM), while SH-SY5Y cells were cultured in DMEM/F12. THP1 cells were cultured in RPMI-1640. Media were supplemented with 10% foetal bovine serum and 1% penicillin–streptomycin with exception of RPMI-1640 which was supplemented with gentamicin (25 μg/mL). Cells were routinely passaged at ∼70–80% confluency using trypsin–EDTA.

### Antisense oligonucleotides

ASOs were designed as 16 nucleotide-long gapmers, with three LNA modified nucleotides at each of the flanks, a DNA gap of 10 nucleotides and a full phosphorothioate backbone. Sequences have a perfect complementarity binding site exclusively in the targeted genes (ENSEMBL_HAVANA genes, version 104^76^).

### ASO transfection and carrier-free uptake

Unless otherwise stated, ASOs were transfected at a final concentration of 20 nM using OPTI-MEM serum-free medium and SilentFect reagent (BioRad) according to the manufacturer’s protocol. For carrier-free uptake (gymnosis) ASOs were added directly to the medium at a final concentration of 5 μM for 7 days.

### CRISPR–Cas9–mediated biallelic dTAG tagging

Biallelic C-terminal dTAG knock-in cell lines were generated using CRISPR–Cas9–mediated genome editing. sgRNAs targeting the region upstream of the endogenous stop codon were designed, and donor templates encoding the dTAG sequence were provided for homology-directed repair. Two donor constructs carrying distinct selection markers (blasticidin or hygromycin resistance) were used to enable tagging of both alleles. Cells were co-transfected with Cas9/sgRNA and donor templates, followed by dual antibiotic selection. Correct biallelic integration was verified by PCR and sequencing, and expression of dTAG-tagged proteins was confirmed by immunoblotting.

### dTAG^V^1 and JTE-607 treatment

dTAG^V^1 (Biotechne) and JTE-607 (Sigma) were dissolved in DMSO to a final concentration of 1 mM and 5 mM, respectively. Cells were supplemented with dTAG^V^1 in the growth media at the final concentration of 1 μM for 24 hours. JTE-607 treatment was performed for 2 hours with JTE-607 added to the growth media at a final concentration of 5 μM. In both cases respective DMSO amount was added to control cells.

### RNAi treatments

Transfections were performed with siLentFect lipid reagent (BioRad) according to the manufacturer’s instructions, with siRNAs at a final concentration of 20 nM. siRNAs used were ON-TARGETplus SMARTpool (Horizon).

### Total RNA extraction

RNA isolation was performed using RNeasy kit (Qiagen). Cells were washed with ice-cold PBS and immediately solubilised in the RLT buffer included in the kit. The next steps were performed according to the kit manual, including on-column DNA digestion using DNase I module (Qiagen).

### chrRNA isolation

Isolation was performed as previously^77^. Collected cells were incubated for 5 min on ice in the Cytoplasmic Lysis Buffer, after which they were layered on 500 µl of Sucrose Buffer (10 mM Tris-HCl pH 7, 150 mM NaCl, 25% Sucrose, 50 U RiboLock). Nuclei were collected by centrifugation at 16000 g for 10 min at 4 °C. The supernatant containing the cytoplasmic fraction was then removed, and nuclei were washed with Nuclei Wash Buffer (PBS supplemented with 0.1% Triton X-100, 1 mM EDTA, 50 U RiboLock) at 1200 g for 1 min at 4 °C. The supernatant was discarded and the nuclei were resuspended in 200 µl of Glycerol Buffer (20 mM Tris-HCl pH 8, 75 mM NaCl, 0.5 mM EDTA, 50% glycerol, 0.85 mM DTT, 50 U RiboLock). Next, 200 µl of Nuclei Lysis Buffer (1% NP-40, 20 mM HEPES pH 7.5, 300 mM NaCl, 1 M Urea, 0.2 mM EDTA, 1 mM DTT, 50 U RiboLock) was mixed with samples by pulsed vortexing for 2 min, and centrifuged at 18500 g for 2 min at 4 °C. The pellet containing chrRNA was resuspended in 200 µl of RNase-free water. The samples were then processed using RNeasy kit (QIAGEN), following the “RNA clean-up” protocol enclosed with the kit, including DNase I on-column treatment. The samples were then quantified using a NanoDrop spectrometer and stored at -80 °C until they were further processed for RT-qPCR or RNA sequencing.

### RT-qPCR

Reverse transcription was performed with Super Script III Reverse Transcriptase (Thermo Fisher) according to the manufacturer’s instructions. Random hexamers and anchored oligo(dT) primers were used for cDNA synthesis. Quantitative PCR (qPCR) was performed using qPCRBIO SyGreen (PCR Biosystems). Reactions were set up according to the manufacturer’s protocol and run on a BioRad real-time PCR system. qPCR data were processed with the ΔΔCt method, with normalization to both GAPDH mRNA levels and the control sample. Primer sequences are shown in Supplementary Table 2.

### PCR analyses

For PCR analyses, HiFi polymerase (PCR Biosystems) was used according to the manufacturer’s protocol. Reactions were run on a BioRad PCR system. PCR products were resolved on 1.5% agarose gel in TAE buffer and visualised using BioRad gel documentation system. Primer sequences are shown in Supplementary Table 2.

### BRB-seq library preparation and sequencing

Total RNA samples (1 μg) were sent to Alithea Genomics SA (Lausanne, Switzerland) for library preparation and sequencing using highly multiplexed 3′-end bulk RNA barcoding and sequencing (MERCURIUSTM BRB-seq service)^33^. The RNA quantity and quality of the RNA was determined using a NanoDrop spectrophotometer (Thermo Fisher Scientific, USA) and the concentration normalized to 100 ng/μl.

#### Library preparation and sequencing

The generation of bulk RNA Barcoding and sequencing (BRB-seq) libraries was performed using the MERCURIUSTM BRB-seq library preparation kit for Illumina and following the manufacturer’s manual (Alithea Genomics, #10813). The library was sequenced on an AVITI from Element Biosciences.

#### Alignment, quantification and data analysis

In this study, we applied STARsolo v2.7.9a^78^ to align and quantify the raw barcodes RNA-seq reads against the GRCh38.104 (human) genome. Utilizing the parameters “--soloUMIdedup NoDedup 1MM_Directional”, and “--quantMode GeneCounts”, we generated raw and UMI-deduplicated count matrices, opting for non-deduplicated counts for subsequent analyses.

### chrRNA-seq library preparation and sequencing

Chromatin RNA samples (500 ng) were submitted to Genomic Technologies facility at the University of Manchester for sequencing. RNA quality and concentrations were evaluated prior to library preparation using standard QC metrics. The RNA was ribo-depleted and libraries were prepared using Illumina Stranded Total RNA Prep with Ribo-Zero library preparation kit and sequenced on the Illumina platform NovaSaq6000 to generate pair-end reads for downstream analysis.

### RNA-seq

RNA was isolated using the MagNAPure 96 system (Roche). The RNA integrity number equivalent (RINe), as assessed with TapeStation High Sensitivity RNA Screen Tape (Agilent), exceeded 8 for all the samples. RNA-seq libraries were prepared with KAPA mRNA HyperPrep Kit (Roche), and sequenced on an Illumina NextSeq 550 system using 75 bp single-end high-output kits.

### Bioinformatics (chrRNA-seq)

Unmapped paired-reads of 59bp were assessed using a quality control pipeline consisting of FastQC v0.12.1 (http://www.bioinformatics.babraham.ac.uk/projects/fastqc/) FastQ Screen v0.15.3 (https://www.bioinformatics.babraham.ac.uk/projects/fastq_screen/), QualiMap v2.2.2-dev^79^ and RSeQC v 4.0.0^80^. The reads were trimmed to remove any adapter or poor quality sequence using Trimmomatic v0.39^81^; the first poor quality base of every read was removed, reads were truncated at a sliding 4bp window (starting 5’) with a mean quality <Q20, and removed if the final length was less than 35 bp. Additional flags included: ’ILLUMINACLIP:./Truseq3-PE-2_Nextera-PE.fa:2:30:10 HEADCROP:1 SLIDINGWINDOW:4:20 MINLEN:35’. The filtered reads were mapped to the human reference sequence analysis set (hg38/Dec. 2013/GRCh38) from the UCSC browser^82^, using STAR v2.7.7a^83^, without gene annotation. Unique, stranded, and concordant paired-end reads ("fragments") were counted into genes using featureCounts from subread v2.0.0, using the parameters -O -s 2 -p -C^84^. The annotation was the comprehensive Gencode v41 gene set^85^. For whole gene analysis the flag -t was set to ’gene’, for the exon only analysis -t was set to ’exon’. To obtain read counts in intron only the exon only reads counts were subtracted from the whole gene read counts using data.frame subtraction in R. Normalisation and differential expression analysis was performed using DESeq2 v1.34.0 on R v4.1.2^86^. A single factor for sample treatment was included in the model. Log fold change shrinkage was applied using the lfcShrink function along with the "apeglm" algorithm. Strand specific fragment coverage profiles were generated from the mapped read output of STAR. Uniquely mapped reads were extracted from the BAM files using samtools view v1.9^87^ applying the flag ’-q 255’. Fragments were split into forward or reverse originating fragments using a script ’split.sh’ from https://www.biostars.org/p/92935/. The files were converted into bedGraph files using genomecov from bedtools v2.25.0^88^. bedGraph counts were scaled by 1/sizeFactor calculated by DESeq2 for the samples in the whole-gene analysis. bigWig files were created using the UCSC Genome Browser tool bedGraphToBigWig.

### *In vitro* deadenylation assay

Briefly, 5 μg of total RNA was mixed with 1 μl 70 mM oligo dT(20), 1 μl of hybridization buffer (10x) and adjusted to a total volume of 10 μL with nuclease-free water. The mix was incubated at 70 °C for 5 min and then slowly cooled to 30 °C. An RNase H buffer (5x) was added together with 2 μl RNase H (Roche) and nuclease-free water to a final volume of 20 μL. Samples were incubated at 30 °C for 30 min and subsequently applied to an RNA purification column (Qiagen). Following washing steps according to the manufacturer’s instructions, RNA was eluted in 25 µl of nuclease-free water and stored at −80 °C until further use.

### *In vitro* RNase H cleavage assay

2 μg *in vitro* T7-synthesised RNA substrate was mixed with 1 µL ASO (2 µM) and 1 μl hybridization buffer in a total volume of 10 μl. Reaction mixtures were incubated at 70 °C for 5 min and then slowly cooled to 37 °C to allow RNA–DNA hybridization. Recombinant RNase H (Roche) was added (0.2 µl per reaction), and reactions were incubated at 37 °C for 20 min. Enzyme activity was terminated by heat inactivation at 90 °C for 2 min, followed by immediate cooling on ice. For PAGE analysis, samples were mixed with 10 µl formamide loading buffer and heated at 70 °C for 5 min prior to electrophoresis. Cleavage products were resolved on 10% TBE denaturing polyacrylamide gel (Invitrogen) run at 180 V for approximately 1 h. Gels were stained in ∼100 mL 0.5x TBE with SYBRSafe for 5–10 min at room temperature, followed by a brief rinse with deionized water and imaged using BioRad gel documentation system.

### Protein extraction

Cells were pelleted down and resuspended in 1 ml extraction buffer (20 mM HEPES-Na, pH 7.4, 0.5% Triton, 600 mM NaCl) supplemented with protease and phosphatase inhibitors (Thermo Fisher). Cell lysate was briefly sonicated and centrifuged for 10 min at 14000 rpm at 4 °C. The supernatant containing protein extract was stored at −80 °C until further use.

### SDS-PAGE and WB analyses

Proteins were separated using the NuPAGE electrophoresis system (Thermo Fisher). Samples were run on 3-8% Tris-Acetate gels. Prestained protein ladder (Thermo Fisher) was used as a marker. Proteins were transferred from the gel to a PVDF membrane by semi-dry transfer with Trans-Blot Turbo Transfer System (Bio-Rad) for 10 min at 25 V. After transfer, membranes were blocked in 5% milk dissolved in PBST buffer supplemented with 0.05% Tween 20. Next, membranes were incubated overnight at 4 °C or for 1 h at room temperature with dedicated primary antibodies (ab178686, ab229078, ab72181, ab176802, 10077-1-AP, AM4300, F3165) followed by incubation with the relevant IRDye secondary antibodies (LICOrbio). Detection was performed using the LICORbio Odyssey Classic Imager.

### Data and code availability

The datasets generated during this study are available at ENA. Accession numbers: PRJEB107235 and PRJEB107065.

## REFERENCES

1. Li, Z., and Rana, T.M. (2014). Therapeutic targeting of microRNAs: current status and future challenges. Nat Rev Drug Discov 13, 622–638. 10.1038/nrd4359.

2. Egli, M., and Manoharan, M. (2023). Chemistry, structure and function of approved oligonucleotide therapeutics. Nucleic Acids Res 51, 2529–2573. 10.1093/nar/gkad067.

3. Havens, M.A., and Hastings, M.L. (2016). Splice-switching antisense oligonucleotides as therapeutic drugs. Nucleic Acids Res 44, 6549–6563. 10.1093/nar/gkw533.

4. Bennett, C.F., Baker, B.F., Pham, N., Swayze, E., and Geary, R.S. (2017). Pharmacology of Antisense Drugs. Annu Rev Pharmacol Toxicol 57, 81–105. 10.1146/annurev-pharmtox-010716-104846.

5. ten Asbroek, A.L., van Groenigen, M., Nooij, M., and Baas, F. (2002). The involvement of human ribonucleases H1 and H2 in the variation of response of cells to antisense phosphorothioate oligonucleotides. Eur J Biochem 269, 583–592. 10.1046/j.0014-2956.2001.02686.x.

6. Wu, H., Lima, W.F., Zhang, H., Fan, A., Sun, H., and Crooke, S.T. (2004). Determination of the role of the human RNase H1 in the pharmacology of DNA-like antisense drugs. J Biol Chem 279, 17181–17189. 10.1074/jbc.M311683200.

7. Monia, B.P., Lesnik, E.A., Gonzalez, C., Lima, W.F., McGee, D., Guinosso, C.J., Kawasaki, A.M., Cook, P.D., and Freier, S.M. (1993). Evaluation of 2’-modified oligonucleotides containing 2’-deoxy gaps as antisense inhibitors of gene expression. J Biol Chem 268, 14514–14522.

8. Lima, W.F., De Hoyos, C.L., Liang, X.H., and Crooke, S.T. (2016). RNA cleavage products generated by antisense oligonucleotides and siRNAs are processed by the RNA surveillance machinery. Nucleic Acids Res 44, 3351–3363. 10.1093/nar/gkw065.

9. Khvorova, A., and Watts, J.K. (2017). The chemical evolution of oligonucleotide therapies of clinical utility. Nat Biotechnol 35, 238–248. 10.1038/nbt.3765.

10. Roberts, T.C., Langer, R., and Wood, M.J.A. (2020). Advances in oligonucleotide drug delivery. Nat Rev Drug Discov 19, 673–694. 10.1038/s41573-020-0075-7.

11. Scoles, D.R., Minikel, E.V., and Pulst, S.M. (2019). Antisense oligonucleotides: A primer. Neurol Genet 5, e323. 10.1212/NXG.0000000000000323.

12. Seth, P.P., Siwkowski, A., Allerson, C.R., Vasquez, G., Lee, S., Prakash, T.P., Wancewicz, E.V., Witchell, D., and Swayze, E.E. (2009). Short antisense oligonucleotides with novel 2’-4’ conformationaly restricted nucleoside analogues show improved potency without increased toxicity in animals. J Med Chem 52, 10–13.10.1021/jm801294h.

13. Swayze, E.E., Siwkowski, A.M., Wancewicz, E.V., Migawa, M.T., Wyrzykiewicz, T.K., Hung, G., Monia, B.P., and Bennett, C.F. (2007). Antisense oligonucleotides containing locked nucleic acid improve potency but cause significant hepatotoxicity in animals. Nucleic Acids Res 35, 687–700. 10.1093/nar/gkl1071.

14. Shen, W., De Hoyos, C.L., Migawa, M.T., Vickers, T.A., Sun, H., Low, A., Bell, T.A., 3rd, Rahdar, M., Mukhopadhyay, S., Hart, C.E., et al. (2019). Chemical modification of PS-ASO therapeutics reduces cellular protein-binding and improves the therapeutic index. Nat Biotechnol 37, 640–650. 10.1038/s41587-019-0106-2.

15. Lai, F., Damle, S.S., Ling, K.K., and Rigo, F. (2020). Directed RNase H Cleavage of Nascent Transcripts Causes Transcription Termination. Mol Cell 77, 1032–1043 e1034. 10.1016/j.molcel.2019.12.029.

16. Lee, J.S., and Mendell, J.T. (2020). Antisense-Mediated Transcript Knockdown Triggers Premature Transcription Termination. Mol Cell 77, 1044–1054 e1043. 10.1016/j.molcel.2019.12.011.

17. Herzel, L., Ottoz, D.S.M., Alpert, T., and Neugebauer, K.M. (2017). Splicing and transcription touch base: co-transcriptional spliceosome assembly and function. Nat Rev Mol Cell Biol 18, 637–650. 10.1038/nrm.2017.63.

18. Oesterreich, F.C., Herzel, L., Straube, K., Hujer, K., Howard, J., and Neugebauer, K.M. (2016). Splicing of Nascent RNA Coincides with Intron Exit from RNA Polymerase II. Cell 165, 372–381. 10.1016/j.cell.2016.02.045.

19. Reimer, K.A., Mimoso, C.A., Adelman, K., and Neugebauer, K.M. (2021). Co-transcriptional splicing regulates 3’ end cleavage during mammalian erythropoiesis. Mol Cell 81, 998–1012 e1017. 10.1016/j.molcel.2020.12.018.

20. Schwalb, B., Michel, M., Zacher, B., Fruhauf, K., Demel, C., Tresch, A., Gagneur, J., and Cramer, P. (2016). TT-seq maps the human transient transcriptome. Science 352, 1225–1228. 10.1126/science.aad9841.

21. Rodriguez-Molina, J.B., West, S., and Passmore, L.A. (2023). Knowing when to stop: Transcription termination on protein-coding genes by eukaryotic RNAPII. Mol Cell 83, 404–415. 10.1016/j.molcel.2022.12.021.

22. Mandel, C.R., Kaneko, S., Zhang, H., Gebauer, D., Vethantham, V., Manley, J.L., and Tong, L. (2006). Polyadenylation factor CPSF-73 is the pre-mRNA 3’-end-processing endonuclease. Nature 444, 953–956. 10.1038/nature05363.

23. Ryan, K., Calvo, O., and Manley, J.L. (2004). Evidence that polyadenylation factor CPSF-73 is the mRNA 3’ processing endonuclease. RNA 10, 565–573. 10.1261/rna.5214404.

24. Rodriguez-Molina, J.B., and Turtola, M. (2023). Birth of a poly(A) tail: mechanisms and control of mRNA polyadenylation. FEBS Open Bio 13, 1140–1153. 10.1002/2211-5463.13528.

25. Proudfoot, N.J. (2011). Ending the message: poly(A) signals then and now. Genes Dev 25, 1770–1782. 10.1101/gad.17268411.

26. Rambout, X., and Maquat, L.E. (2024). Nuclear mRNA decay: regulatory networks that control gene expression. Nat Rev Genet 25, 679–697. 10.1038/s41576-024-00712-2.

27. Cortazar, M.A., Sheridan, R.M., Erickson, B., Fong, N., Glover-Cutter, K., Brannan, K., and Bentley, D.L. (2019). Control of RNA Pol II Speed by PNUTS-PP1 and Spt5 Dephosphorylation Facilitates Termination by a "Sitting Duck Torpedo" Mechanism. Mol Cell 76, 896–908 e894. 10.1016/j.molcel.2019.09.031.

28. Eaton, J.D., Francis, L., Davidson, L., and West, S. (2020). A unified allosteric/torpedo mechanism for transcriptional termination on human protein-coding genes. Genes Dev 34, 132–145. 10.1101/gad.332833.119.

29. Marasco, L.E., Dujardin, G., Sousa-Luis, R., Liu, Y.H., Stigliano, J.N., Nomakuchi, T., Proudfoot, N.J., Krainer, A.R., and Kornblihtt, A.R. (2022). Counteracting chromatin effects of a splicing-correcting antisense oligonucleotide improves its therapeutic efficacy in spinal muscular atrophy. Cell 185, 2057–2070 e2015. 10.1016/j.cell.2022.04.031.

30. Herrmann, C.J., Schmidt, R., Kanitz, A., Artimo, P., Gruber, A.J., and Zavolan, M. (2020). PolyASite 2.0: a consolidated atlas of polyadenylation sites from 3’ end sequencing. Nucleic Acids Res 48, D174–D179. 10.1093/nar/gkz918.

31. Zhang, H., Hu, J., Recce, M., and Tian, B. (2005). PolyA_DB: a database for mammalian mRNA polyadenylation. Nucleic Acids Res 33, D116–120. 10.1093/nar/gki055.

32. Wang, R., Nambiar, R., Zheng, D., and Tian, B. (2018). PolyA_DB 3 catalogs cleavage and polyadenylation sites identified by deep sequencing in multiple genomes. Nucleic Acids Res 46, D315–D319. 10.1093/nar/gkx1000.

33. Alpern, D., Gardeux, V., Russeil, J., Mangeat, B., Meireles-Filho, A.C.A., Breysse, R., Hacker, D., and Deplancke, B. (2019). BRB-seq: ultra-affordable high-throughput transcriptomics enabled by bulk RNA barcoding and sequencing. Genome Biol 20, 71. 10.1186/s13059-019-1671-x.

34. Mayer, A., di Iulio, J., Maleri, S., Eser, U., Vierstra, J., Reynolds, A., Sandstrom, R., Stamatoyannopoulos, J.A., and Churchman, L.S. (2015). Native elongating transcript sequencing reveals human transcriptional activity at nucleotide resolution. Cell 161, 541–554. 10.1016/j.cell.2015.03.010.

35. Robinson, J.T., Thorvaldsdottir, H., Winckler, W., Guttman, M., Lander, E.S., Getz, G., and Mesirov, J.P. (2011). Integrative genomics viewer. Nat Biotechnol 29, 24–26. 10.1038/nbt.1754.

36. Gagnon, K.T., and Corey, D.R. (2019). Guidelines for Experiments Using Antisense Oligonucleotides and Double-Stranded RNAs. Nucleic Acid Ther 29, 116–122. 10.1089/nat.2018.0772.

37. Dhanasekaran, R., Deutzmann, A., Mahauad-Fernandez, W.D., Hansen, A.S., Gouw, A.M., and Felsher, D.W. (2022). The MYC oncogene - the grand orchestrator of cancer growth and immune evasion. Nat Rev Clin Oncol 19, 23–36. 10.1038/s41571-021-00549-2.

38. Chan, A.M., Takai, S., Yamada, K., and Miki, T. (1996). Isolation of a novel oncogene, NET1, from neuroepithelioma cells by expression cDNA cloning. Oncogene 12, 1259–1266.

39. Kakegawa, J., Sakane, N., Suzuki, K., and Yoshida, T. (2019). JTE-607, a multiple cytokine production inhibitor, targets CPSF3 and inhibits pre-mRNA processing. Biochem Biophys Res Commun 518, 32–37. 10.1016/j.bbrc.2019.08.004.

40. Ross, N.T., Lohmann, F., Carbonneau, S., Fazal, A., Weihofen, W.A., Gleim, S., Salcius, M., Sigoillot, F., Henault, M., Carl, S.H., et al. (2020). CPSF3-dependent pre-mRNA processing as a druggable node in AML and Ewing’s sarcoma. Nat Chem Biol 16, 50–59. 10.1038/s41589-019-0424-1.

41. Liu, L., Yu, A.M., Wang, X., Soles, L.V., Teng, X., Chen, Y., Yoon, Y., Sarkan, K.S.K., Valdez, M.C., Linder, J., et al. (2023). The anticancer compound JTE-607 reveals hidden sequence specificity of the mRNA 3’ processing machinery. Nat Struct Mol Biol 30, 1947–1957. 10.1038/s41594-023-01161-x.

42. Tomecki, R., Kristiansen, M.S., Lykke-Andersen, S., Chlebowski, A., Larsen, K.M., Szczesny, R.J., Drazkowska, K., Pastula, A., Andersen, J.S., Stepien, P.P., et al. (2010). The human core exosome interacts with differentially localized processive RNases: hDIS3 and hDIS3L. EMBO J 29, 2342–2357. 10.1038/emboj.2010.121.

43. Nabet, B., Roberts, J.M., Buckley, D.L., Paulk, J., Dastjerdi, S., Yang, A., Leggett, A.L., Erb, M.A., Lawlor, M.A., Souza, A., et al. (2018). The dTAG system for immediate and target-specific protein degradation. Nat Chem Biol 14, 431–441. 10.1038/s41589-018-0021-8.

44. Preker, P., Nielsen, J., Kammler, S., Lykke-Andersen, S., Christensen, M.S., Mapendano, C.K., Schierup, M.H., and Jensen, T.H. (2008). RNA exosome depletion reveals transcription upstream of active human promoters. Science 322, 1851–1854. 10.1126/science.1164096.

45. West, S., Gromak, N., Norbury, C.J., and Proudfoot, N.J. (2006). Adenylation and exosome-mediated degradation of cotranscriptionally cleaved pre-messenger RNA in human cells. Mol Cell 21, 437–443. 10.1016/j.molcel.2005.12.008.

46. Wu, G., Schmid, M., Rib, L., Polak, P., Meola, N., Sandelin, A., and Jensen, T.H. (2020). A Two-Layered Targeting Mechanism Underlies Nuclear RNA Sorting by the Human Exosome. Cell Rep 30, 2387–2401 e2385. 10.1016/j.celrep.2020.01.068.

47. Buntz, A., Killian, T., Schmid, D., Seul, H., Brinkmann, U., Ravn, J., Lindholm, M., Knoetgen, H., Haucke, V., and Mundigl, O. (2019). Quantitative fluorescence imaging determines the absolute number of locked nucleic acid oligonucleotides needed for suppression of target gene expression. Nucleic Acids Res 47, 953–969. 10.1093/nar/gky1158.

48. Kielpinski, L.J., Funder, E.D., Schmidt, S., and Hagedorn, P.H. (2021). Characterization of Escherichia coli RNase H Discrimination of DNA Phosphorothioate Stereoisomers. Nucleic Acid Ther 31, 383–391. 10.1089/nat.2021.0055.

49. Hansen, H.F., Albaek, N., Hansen, B.R., Shim, I., Bohr, H., and Koch, T. (2021). In vivo uptake of antisense oligonucleotide drugs predicted by ab initio quantum mechanical calculations. Sci Rep 11, 6321. 10.1038/s41598-021-85453-6.

50. Papargyri, N., Pontoppidan, M., Andersen, M.R., Koch, T., and Hagedorn, P.H. (2020). Chemical Diversity of Locked Nucleic Acid-Modified Antisense Oligonucleotides Allows Optimization of Pharmaceutical Properties. Mol Ther Nucleic Acids 19, 706–717. 10.1016/j.omtn.2019.12.011.

51. Pendergraff, H., Schmidt, S., Vikesa, J., Weile, C., Overup, C., M, W.L., and Koch, T. (2020). Nuclear and Cytoplasmatic Quantification of Unconjugated, Label-Free Locked Nucleic Acid Oligonucleotides. Nucleic Acid Ther 30, 4–13. 10.1089/nat.2019.0810.

52. Andersson, P., Burel, S.A., Estrella, H., Foy, J., Hagedorn, P.H., Harper, T.A., Jr., Henry, S.P., Hoflack, J.C., Holgersen, E.M., Levin, A.A., et al. (2025). Assessing Hybridization-Dependent Off-Target Risk for Therapeutic Oligonucleotides: Updated Industry Recommendations. Nucleic Acid Ther 35, 16–33. 10.1089/nat.2024.0072.

53. Helm, J., Schols, L., and Hauser, S. (2022). Towards Personalized Allele-Specific Antisense Oligonucleotide Therapies for Toxic Gain-of-Function Neurodegenerative Diseases. Pharmaceutics 14. 10.3390/pharmaceutics14081708.

54. Ziegler, A., Carroll, J., Bain, J.M., Sands, T.T., Fee, R.J., Uher, D., Kanner, C.H., Montes, J., Glass, S., Douville, J., et al. (2024). Antisense oligonucleotide therapy in an individual with KIF1A-associated neurological disorder. Nat Med 30, 2782–2786. 10.1038/s41591-024-03197-y.

55. Anderson, B.R., Jensen, M.L., Hagedorn, P.H., Little, S.C., Olson, R.E., Ammar, R., Kienzle, B., Thompson, J., McDonald, I., Mercer, S., et al. (2020). Allele-Selective Knockdown of MYH7 Using Antisense Oligonucleotides. Mol Ther Nucleic Acids 19, 1290–1298. 10.1016/j.omtn.2020.01.012.

56. Crooke, S.T. (2024). Addressing the Challenges of Treating Patients with Heterozygous Gain of Function Mutations. Nucleic Acid Ther 34, 273–275. 10.1089/nat.2024.0060.

57. Liang, X.H., Vickers, T.A., Guo, S., and Crooke, S.T. (2011). Efficient and specific knockdown of small non-coding RNAs in mammalian cells and in mice. Nucleic Acids Res 39, e13. 10.1093/nar/gkq1121.

58. Tripathi, V., Ellis, J.D., Shen, Z., Song, D.Y., Pan, Q., Watt, A.T., Freier, S.M., Bennett, C.F., Sharma, A., Bubulya, P.A., et al. (2010). The nuclear-retained noncoding RNA MALAT1 regulates alternative splicing by modulating SR splicing factor phosphorylation. Mol Cell 39, 925–938. 10.1016/j.molcel.2010.08.011.

59. Shen, W., Liang, X.H., and Crooke, S.T. (2014). Phosphorothioate oligonucleotides can displace NEAT1 RNA and form nuclear paraspeckle-like structures. Nucleic Acids Res 42, 8648–8662. 10.1093/nar/gku579.

60. Li, Y., Garcia, G., Jr., Arumugaswami, V., and Guo, F. (2021). Structure-based design of antisense oligonucleotides that inhibit SARS-CoV-2 replication. bioRxiv. 10.1101/2021.08.23.457434.

61. Pollak, A.J., Hickman, J.H., Liang, X.H., and Crooke, S.T. (2020). Gapmer Antisense Oligonucleotides Targeting 5S Ribosomal RNA Can Reduce Mature 5S Ribosomal RNA by Two Mechanisms. Nucleic Acid Ther 30, 312–324. 10.1089/nat.2020.0864.

62. Horberg, J., Carlesso, A., and Reymer, A. (2024). Mechanistic insights into ASO-RNA complexation: Advancing antisense oligonucleotide design strategies. Mol Ther Nucleic Acids 35, 102351. 10.1016/j.omtn.2024.102351.

63. Chao, L.C., Jamil, A., Kim, S.J., Huang, L., and Martinson, H.G. (1999). Assembly of the cleavage and polyadenylation apparatus requires about 10 seconds in vivo and is faster for strong than for weak poly(A) sites. Mol Cell Biol 19, 5588–5600. 10.1128/MCB.19.8.5588.

64. Hyjek, M., Figiel, M., and Nowotny, M. (2019). RNases H: Structure and mechanism. DNA Repair (Amst) 84, 102672. 10.1016/j.dnarep.2019.102672.

65. Skourti-Stathaki, K., and Proudfoot, N.J. (2014). A double-edged sword: R loops as threats to genome integrity and powerful regulators of gene expression. Genes Dev 28, 1384–1396. 10.1101/gad.242990.114.

66. Petermann, E., Lan, L., and Zou, L. (2022). Sources, resolution and physiological relevance of R-loops and RNA-DNA hybrids. Nat Rev Mol Cell Biol 23, 521–540. 10.1038/s41580-022-00474-x.

67. Meola, N., Domanski, M., Karadoulama, E., Chen, Y., Gentil, C., Pultz, D., Vitting-Seerup, K., Lykke-Andersen, S., Andersen, J.S., Sandelin, A., and Jensen, T.H. (2016). Identification of a Nuclear Exosome Decay Pathway for Processed Transcripts. Mol Cell 64, 520–533. 10.1016/j.molcel.2016.09.025.

68. Lubas, M., Christensen, M.S., Kristiansen, M.S., Domanski, M., Falkenby, L.G., Lykke-Andersen, S., Andersen, J.S., Dziembowski, A., and Jensen, T.H. (2011). Interaction profiling identifies the human nuclear exosome targeting complex. Mol Cell 43, 624–637. 10.1016/j.molcel.2011.06.028.

69. Warkocki, Z., Liudkovska, V., Gewartowska, O., Mroczek, S., and Dziembowski, A. (2018). Terminal nucleotidyl transferases (TENTs) in mammalian RNA metabolism. Philos Trans R Soc Lond B Biol Sci 373. 10.1098/rstb.2018.0162.

70. Singh, N.N., Luo, D., and Singh, R.N. (2018). Pre-mRNA Splicing Modulation by Antisense Oligonucleotides. Methods Mol Biol 1828, 415–437. 10.1007/978-1-4939-8651-4_26.

71. Tian, B., and Graber, J.H. (2012). Signals for pre-mRNA cleavage and polyadenylation. Wiley Interdiscip Rev RNA 3, 385–396. 10.1002/wrna.116.

72. Grzechnik, P., and Mischo, H.E. (2025). Fateful Decisions of Where to Cut the Line: Pathology Associated with Aberrant 3’ End Processing and Transcription Termination. J Mol Biol 437, 168802. 10.1016/j.jmb.2024.168802.

73. Hill, C.H., Boreikaite, V., Kumar, A., Casanal, A., Kubik, P., Degliesposti, G., Maslen, S., Mariani, A., von Loeffelholz, O., Girbig, M., et al. (2019). Activation of the Endonuclease that Defines mRNA 3’ Ends Requires Incorporation into an 8-Subunit Core Cleavage and Polyadenylation Factor Complex. Mol Cell 73, 1217–1231 e1211. 10.1016/j.molcel.2018.12.023.

74. Torres-Ulloa, L., Calvo-Roitberg, E., and Pai, A.A. (2024). Genome-wide kinetic profiling of pre-mRNA 3’ end cleavage. RNA 30, 256–270. 10.1261/rna.079783.123.

75. Baejen, C., Andreani, J., Torkler, P., Battaglia, S., Schwalb, B., Lidschreiber, M., Maier, K.C., Boltendahl, A., Rus, P., Esslinger, S., et al. (2017). Genome-wide Analysis of RNA Polymerase II Termination at Protein-Coding Genes. Mol Cell 66, 38–49 e36. 10.1016/j.molcel.2017.02.009.

76. Dyer, S.C., Austine-Orimoloye, O., Azov, A.G., Barba, M., Barnes, I., Barrera-Enriquez, V.P., Becker, A., Bennett, R., Beracochea, M., Berry, A., et al. (2025). Ensembl 2025. Nucleic Acids Res 53, D948–D957. 10.1093/nar/gkae1071.

77. Sledziowska, M., Winczura, K., Jones, M., Almaghrabi, R., Mischo, H., Hebenstreit, D., Garcia, P., and Grzechnik, P. (2023). Non-coding RNAs associated with Prader-Willi syndrome regulate transcription of neurodevelopmental genes in human induced pluripotent stem cells. Hum Mol Genet 32, 608–620. 10.1093/hmg/ddac228.

78. Kaminow, B., Yunusov, D., and Dobin, A. (2021). STARsolo: accurate, fast and versatile mapping/quantification of single-cell and single-nucleus RNA-seq data. bioRxiv, 2021.2005.2005.442755. 10.1101/2021.05.05.442755.

79. Okonechnikov, K., Conesa, A., and Garcia-Alcalde, F. (2016). Qualimap 2: advanced multi-sample quality control for high-throughput sequencing data. Bioinformatics 32, 292–294. 10.1093/bioinformatics/btv566.

80. Wang, L., Wang, S., and Li, W. (2012). RSeQC: quality control of RNA-seq experiments. Bioinformatics 28, 2184–2185. 10.1093/bioinformatics/bts356.

81. Bolger, A.M., Lohse, M., and Usadel, B. (2014). Trimmomatic: a flexible trimmer for Illumina sequence data. Bioinformatics 30, 2114–2120. 10.1093/bioinformatics/btu170.

82. Kent, W.J., Sugnet, C.W., Furey, T.S., Roskin, K.M., Pringle, T.H., Zahler, A.M., and Haussler, D. (2002). The human genome browser at UCSC. Genome Res 12, 996–1006. 10.1101/gr.229102.

83. Dobin, A., Davis, C.A., Schlesinger, F., Drenkow, J., Zaleski, C., Jha, S., Batut, P., Chaisson, M., and Gingeras, T.R. (2013). STAR: ultrafast universal RNA-seq aligner. Bioinformatics 29, 15–21. 10.1093/bioinformatics/bts635.

84. Liao, Y., Smyth, G.K., and Shi, W. (2014). featureCounts: an efficient general purpose program for assigning sequence reads to genomic features. Bioinformatics 30, 923–930. 10.1093/bioinformatics/btt656.

85. Harrow, J., Frankish, A., Gonzalez, J.M., Tapanari, E., Diekhans, M., Kokocinski, F., Aken, B.L., Barrell, D., Zadissa, A., Searle, S., et al. (2012). GENCODE: the reference human genome annotation for The ENCODE Project. Genome Res 22, 1760–1774. 10.1101/gr.135350.111.

86. Love, M.I., Huber, W., and Anders, S. (2014). Moderated estimation of fold change and dispersion for RNA-seq data with DESeq2. Genome Biol 15, 550. 10.1186/s13059-014-0550-8.

87. Li, H., Handsaker, B., Wysoker, A., Fennell, T., Ruan, J., Homer, N., Marth, G., Abecasis, G., Durbin, R., and Genome Project Data Processing, S. (2009). The Sequence Alignment/Map format and SAMtools. Bioinformatics 25, 2078–2079. 10.1093/bioinformatics/btp352.

88. Quinlan, A.R., and Hall, I.M. (2010). BEDTools: a flexible suite of utilities for comparing genomic features. Bioinformatics 26, 841–842. 10.1093/bioinformatics/btq033.

